# Rat and rabbit whole embryo culture as a new approach method for unlabeled therapeutic antisense oligonucleotide hazard identification with no requirement for microinjection or assisted transfection

**DOI:** 10.64898/2025.12.23.692174

**Authors:** Sara M. Bender, Sharon Chapman, Sreenivas Nannapaneni, Joel D. Parry, Joyce Rendemonti, Stanley Robinson, Bradley Spencer-Dene, Dinesh Stanislaus

## Abstract

The developmental hazard screening of oligonucleotide therapeutics (ONTs) presents challenges due to their frequent lack of pharmacology in nonclinical species and embryofetal exposure is presumed to be limited *in vivo*. This study demonstrates that direct culture in media containing mipomersen, a 2’-O-methoxyethyl phosphorothioated antisense oligonucleotide (ASO), results in dose-responsive morphological changes in rat and rabbit whole embryo culture (WEC). Automated miRNAscope *in situ* hybridization was used to confirm dose-dependent embryonic exposure and visualize the distribution pattern of mipomersen in the embryo and extra-embryonic membranes, suggesting that ONTs may enter the umbilical vasculature and pass into embryo circulation. Neither microinjections nor assisted transfections were required to achieve embryonic exposure. These findings support the utility of WEC as a new approach method (NAM) for developmental hazard identification of ONTs. WEC could complement or partially replace *in vivo* studies, reducing animal use and required test material amounts, while enabling robust developmental hazard identification for ONTs. This work informs future safety assessment strategies and regulatory guidance for ONTs.

## Introduction

The emergence of oligonucleotide therapeutics (ONTs) has challenged the pharmaceutical industry to develop new strategies and standard practices for nonclinical safety assessment of this modality, which has characteristics of both small molecules and biologics. Biodistribution of ONTs in developmental and reproductive toxicity (DART) assessments is an important consideration, as nearly all marketed ONTs have been detected in placentas of nonclinical species but generally have not been detectable in fetal liver, kidney or homogenized whole fetus^1^. However, two marketed GalNAc-siRNAs, LEQVIO (inclisiran sodium) and RIVFLOZA (nedosiran sodium), have been detected in fetal plasma^1^. Depending on ONT-specific dosing strategies and *in vivo* DART study design considerations, it may not be feasible to achieve meaningful ONT exposure throughout the entire period of organogenesis.

Whole embryo culture (WEC) is an *ex vivo* hazard identification screening assay and is considered a new approach method (NAM). WEC allows rodent or rabbit embryos to be directly exposed to the test item during an early critical period of major organogenesis, offering the opportunity to perform developmental hazard assessment for ONTs that may not achieve meaningful fetal exposure during *in vivo* DART studies or when surrogate molecule use is necessary. It was previously thought that microinjection into the amniotic cavity, the embryo itself or the largest yolk sac blood vessels was necessary to ensure exposure to the embryo due to the larger size of ONTs^2^. Previous studies have successfully evaluated ONTs in mouse WEC using a microinjection approach^3–6^. In contrast, WEC studies evaluating small molecules are regularly performed, as they are presumed to be small enough to diffuse into the yolk sac cavity and be absorbed into the embryo. While it is unlikely that ONTs could diffuse through the yolk sac, endocytosis could potentially allow for their uptake resulting in exposure to embryos cultured in media containing ONTs,^7^ given gymnotic exposure was successful. This approach is much simpler in terms of technical execution as compared to the microinjection approach and prevents experimental artifacts that may interfere with interpretation. Demonstration that microinjection is not necessary for ONTs would greatly increase the utility of WEC studies for developmental toxicity hazard assessments.

KYNAMRO (mipomersen), a 20-nucleotide long, 2’-O-methoxyethyl phosphorothioated ASO (2’-MOE ASO), was used to investigate the utility of rat and rabbit WEC to assess ONTs using unassisted transfection. Mipomersen was approved by the FDA to treat homozygous familial hypercholesterolemia and is an inhibitor of apolipoprotein B-100 synthesis, a protein essential for the production of low-density lipoprotein (LDL), although it was later withdrawn from the market due to unacceptable risk of hepatotoxicity^8^. Mipomersen is not pharmacologically active in rats or rabbits^9^.

For the first time, we show concentration dependent embryonic exposure to an ONT following culture in media containing 2’-MOE ASO (mipomersen) alone, with no microinjection, for both rat and rabbits on gestation day (GD) 11. Exposure was confirmed via off-target dose-responsive effects on yolk sac and/or embryo parameters and automated miRNAscope *in situ* hybridization showing distribution of mipomersen in embryos and yolk sac umbilical vasculature at the point of connection with the embryo.

## Materials & Methods

### Rat Husbandry and WEC

Rat embryos were cultured according to established techniques^10^. Untreated time-mated female Sprague–Dawley rats (approximately 10 to 12 weeks of age, supplied by Charles River, Laboratories, Inc., Raleigh, NC) were used for this study. Pregnant dams were approximately Gestation Day (GD) 5 upon arrival (day of mating is designated as GD 0).

Mated female rats were housed in Allentown NexGen reusable individually ventilated caging (IVC) system with clear plastic cages containing Alpha-dri™ bedding (Shepherd Specialty Papers, Inc., Kalamazoo, MI) in a controlled environment (68°F-79°F; 30% to 70% relative humidity) with an approximate 12-hour light/12-hour dark cycle. Food (Certified Rodent Diet #5001, PMI Nutrition International Brentwood, MO) was provided ad libitum.

Filtered tap water (supplied and periodically tested by American Water) was available ad libitum from an automatic watering system. Rat enrichment (plastic or cardboard hut or tube, wood gnawing block) was provided and documented. Available information indicated that no substance was present in the diet, drinking water, enrichment or bedding at a concentration likely to influence the outcome of this study.

Dams were anesthetized to a deep plane of anesthesia with isoflurane and euthanized via exsanguination in the afternoon on GD 9. Death was ensured by thoracotomy as a secondary form of euthanasia. Embryos were dissected out of the uterus for subsequent whole embryo culture. Using a dissecting microscope, the Reichert’s membrane was removed and embryos at the appropriate stage of development (i.e., late headfold stage and 1 or 2 somite pair) were chosen for the experiment^11^. Selected embryos were placed into pre-warmed, sterile bottles containing approximately 2.0 mL culture media (70% heat-inactivated rat WEC serum, 30% Tyrode’s solution, 35 μg/ml streptomycin) as well as either vehicle control (0.04% PBS) or 1, 10, 50 or 100 µM mipomersen (WuXi). The culture bottles were maintained at 37°C ± 0.5 °C in an incubator (Cullum Starr Precision Engineering Limited, Cambridge, England) for approximately 40 hours. The bottles rotated at a rate of approximately 30 rotations per minute and were continuously gassed using an intermediate low flow regulator set at a final flow rate of approximately 30 cc/min with scheduled, increasing oxygen concentrations from 5 to 10 to 20% O_2_, (with a constant 5% of CO_2_ and the balance of nitrogen between 75 and 90%). Ninety-four embryos were individually cultured, 30 vehicle control and 16 treated per group. Embryos were distributed such that each litter was represented in each group and each group was distributed between two or three incubators. At the end of the culture period on GD 11, embryos were examined for viability, morphology, size (crown-rump length), and somite number using a modified version of a Morphological Scoring System^12^. In addition, visceral yolk sacs (VYS) were evaluated for morphology and size (diameter) prior to embryo evaluation.

### Rabbit Husbandry and WEC

Untreated time-mated female New Zealand White rabbits (approximately 6 to 9 months of age, supplied by Envigo Global Services, Inc., Denver, PA) were used for this study. Pregnant does were approximately GD 3 to 5 upon arrival (day of mating designated as GD 0). Mated female rabbits were housed individually in stainless steel Allentown Rabbit Euro Cages with plastic fenestrated flooring in a controlled environment (61°F-72°F; 30% to 70% relative humidity) with an approximate 12-hour light/12-hour dark cycle. Food [Rabbit Chow (HF) #5322, PMI Nutritional International, Brentwood, MO] was provided (approximately 125g daily). Filtered tap water (supplied and periodically tested by American Water) was available ad libitum from an automatic watering system. Animals received appropriate music for a period during the light hours. Rabbit enrichment (plastic resting board with hide, wood gnawing block, stainless steel bell, nylon dumbbells) was provided and documented. Available information indicates that no substance is present in the diet, drinking water, or enrichment at a concentration likely to influence the outcome of this study.

Does were euthanized in the morning on GD 9 via intravenous overdose of Fatal Plus® (pentobarbital sodium) via the marginal ear vein. Death was ensured via exsanguination as a secondary form of euthanasia. Embryos were dissected out of the uterus and the bed of chorionic tissue was removed, such that only the embryo, amnion and attached visceral yolk sac were included for subsequent whole embryo culture. Criteria for placement into culture was as follows: clearly developed eyes, embryo positioned all on the same side of the yolk sac, intact vessel encircling yolk sac, visible blood flow, and lack of abnormalities. Selected embryos were placed into pre-warmed, sterile bottles containing approximately 2.5 mL culture media [75% heat-activated rabbit WEC serum, 25% phosphate buffered saline (PBS), 10 µg/mL streptomycin, 10 µl/mL penicillin, 2 mg/mL glucose] as well as either vehicle control (0.04% PBS) or 1, 10, 50 or 100 µM mipomersen (WuXi). The culture bottles were maintained at 37°C ± 0.5 °C in an incubator (Cullum Starr Precision Engineering Limited, Cambridge, England) for approximately 45 hours. The bottles rotated at a rate of approximately 30 rotations per minute and were continuously gassed using an intermediate low flow regulator set at a final flow rate of approximately 30 cc/min with scheduled, increasing oxygen concentrations from 20 to 95% O_2_, (with a constant 5% of CO_2_ and the balance of nitrogen between 0 and 75%). Ninety-four embryos were individually cultured, 30 vehicle control and 16 treated per group. Embryos were distributed such that each litter was represented in each group, and each group was distributed between two or three incubators. At the end of the culture period on GD 11, embryos were examined for morphology, size (crown-rump length), and somite number using a modified version of a Morphological Scoring System^13,14^. In addition, visceral yolk sacs (VYS) were evaluated for morphology and size (diameter) prior to embryo evaluation.

### Embryo Evaluation of Morphology and Fixation

On GD11, rat and rabbit embryos were dissected from the yolk sac and amnion, which were retained, and evaluated for viability, morphology, size and somite number. Visceral yolk sacs were evaluated for morphology and size. Dead and/or grossly malformed embryos were only evaluated for viability. Embryos found outside the yolk sac following the culture period were not evaluated as these were considered experimental artifacts.

The following regions were evaluated for both rat and rabbit: yolk sac, embryo rotation, somite, caudal, forelimb, hindlimb, neural tube, heart, pharyngeal arches, and craniofacial. Supplemental Tables 2 and 4 list all detailed potential observations for each region. In the heart region, observations of dextrocardia were recorded in the raw data, but not summarized in Table 1, because this finding is considered an artifact of culture and not treatment related.

**Table 1.**

| Rat WEC Morphology Summary |  |  |  |  |  |
| --- | --- | --- | --- | --- | --- |
| Dose Level | 0 μM | 1 μM | 10 μM | 50 μM | 100 μM |
| **Able to Examine** |  |  |  |  |  |
| Yes | 30 | 16 | 16 | 16 | 4 |
| No | 0 | 0 | 0 | 0 | 12 |
| Dead | 0 | 0 | 0 | 0 | 1 |
| Grossly Malformed | 0 | 0 | 0 | 0 | 11 |
| **Caudal** |  |  |  |  |  |
| Total Embryos with Caudal Abnormalities | 0 | 0 | 1* | 8* | 4* |
| % Examined Embryos with Caudal Abnormalities | 0.0% | 0.0% | 6.3% | 50.0% | 100.0% |
| **Craniofacial** |  |  |  |  |  |
| Total Embryos with Craniofacial Abnormalities | 0 | 0 | 0 | 7* | 4* |
| % Examined Embryos with Craniofacial Abnormalities | 0.0% | 0.0% | 0.0% | 43.8% | 100.0% |
| **Forelimb Bud** |  |  |  |  |  |
| Total Embryos with Forelimb Bud Abnormalities | 0 | 0 | 2* | 10* | 4* |
| % Examined Embryos with Forelimb Bud Abnormalities | 0.0% | 0.0% | 12.5% | 62.5% | 100.0% |
| **Hindlimb Bud** |  |  |  |  |  |
| Total Embryos with Hindlimb Bud Abnormalities | 0 | 0 | 0 | 3* | 4* |
| % Examined Embryos with Hindlimb Bud Abnormalities | 0.0% | 0.0% | 0.0% | 18.8% | 100.0% |
| **Heart** |  |  |  |  |  |
| Total Embryos with Heart Abnormalities | 1 | 0 | 1* | 4* | 0*† |
| % Examined Embryos with Heart Abnormalities | 3.3% | 0.0% | 6.3% | 25.0% | 0.0% |
| **Neural Tube** |  |  |  |  |  |
| Total Embryos with Neural Tube Abnormalities | 0 | 0 | 0 | 7* | 4* |
| % Examined Embryos with Neural Tube Abnormalities | 0.0% | 0.0% | 0.0% | 43.8% | 100.0% |
| **Pharyngeal Arches** |  |  |  |  |  |
| Total Embryos with Pharyngeal Arch Abnormalities | 0 | 0 | 0 | 3* | 2* |
| % Examined Embryos with Pharyngeal Arch Abnormalities | 0.0% | 0.0% | 0.0% | 18.8% | 50.0% |
| **Rotation** |  |  |  |  |  |
| Total Embryos with Rotation Abnormalities | 1 | 0 | 0 | 7* | 4* |
| % Examined Embryos with Rotation Abnormalities | 3.3% | 0.0% | 0.0% | 43.8% | 100.0% |
| **Somite Morphology** |  |  |  |  |  |
| Total Embryos with Somite Abnormalities | 0 | 0 | 0 | 1 | 3* |
| % Examined Embryos with Somite Abnormalities | 0.0% | 0.0% | 0.0% | 6.3% | 75.0% |
| **Yolk Sac Morphology** |  |  |  |  |  |
| Total Yolk Sacs with any abnormalities | 0 | 0 | 0 | 2* | 4* |
| % Examined Yolk Sacs with any abnormalities | 0.0% | 0.0% | 0.0% | 12.5% | 100.0% |
| **Rabbit WEC Morphology Summary** |  |  |  |  |  |
| Dose Level | 0 μM | 1 μM | 10 μM | 50 μM | 100 μM |
| **Able to Examine** |  |  |  |  |  |
| Yes | 25 | 12 | 15 | 13 | 15 |
| No | 5 | 4 | 1 | 3 | 1 |
| Dead | 0 | 1 | 0 | 0 | 0 |
| Grossly Malformed | 0 | 0 | 0 | 1 | 1 |
| Outside VYS | 5 | 3 | 1 | 2 | 0 |
| **Caudal** |  |  |  |  |  |
| Total Embryos with Caudal Abnormalities | 1 | 2 | 2 | 1 | 2 |
| % Examined Embryos with Caudal Abnormalities | 4.0% | 16.7% | 13.3% | 7.7% | 13.3% |
| **Craniofacial** |  |  |  |  |  |
| Total Embryos with Craniofacial Abnormalities | 3 | 2 | 2 | 1 | 5* |
| % Examined Embryos with Craniofacial Abnormalities | 12.0% | 16.7% | 13.3% | 7.7% | 33.3% |
| **Forelimb Bud** |  |  |  |  |  |
| Total Embryos with Forelimb Bud Abnormalities | 8 | 4 | 4 | 6 | 5 |
| % Examined Embryos with Forelimb Bud Abnormalities | 32.0% | 33.3% | 26.7% | 46.2% | 33.3% |
| **Hindlimb Bud** |  |  |  |  |  |
| Total Embryos with Hindlimb Bud Abnormalities | 3 | 2 | 2 | 2 | 6* |
| % Examined Embryos with Hindlimb Bud Abnormalities | 12.0% | 16.7% | 13.3% | 15.4% | 40.0% |
| **Heart** |  |  |  |  |  |
| Total Embryos with Heart Abnormalities | 0 | 0 | 2 | 0 | 3* |
| % Examined Embryos with Heart Abnormalities | 0.0% | 0.0% | 13.3% | 0.0% | 20.0% |
| **Neural Tube** |  |  |  |  |  |
| Total Embryos with Neural Tube Abnormalities | 3 | 1 | 2 | 2 | 4* |
| % Examined Embryos with Neural Tube Abnormalities | 12.0% | 8.3% | 13.3% | 15.4% | 26.7 |
| **Pharyngeal Arches** |  |  |  |  |  |
| Total Embryos with Pharyngeal Arch Abnormalities | 0 | 1 | 1 | 1 | 0 |
| % Examined Embryos with Pharyngeal Arch Abnormalities | 0.0% | 8.3% | 6.7% | 7.7% | 0.0% |
| **Rotation** |  |  |  |  |  |
| Total Embryos with Rotation Abnormalities | 5 | 3 | 3 | 3 | 7* |
| % Examined Embryos with Rotation Abnormalities | 20.0% | 25.0% | 20.0% | 23.1% | 46.7% |
| **Somite Morphology** |  |  |  |  |  |
| Total Embryos with Somite Abnormalities | 0 | 1 | 1 | 0 | 3* |
| % Examined Embryos with Somite Abnormalities | 0.0% | 8.3% | 6.7% | 0.0% | 20.0% |
| **Yolk Sac Morphology** |  |  |  |  |  |
| Total Yolk Sacs with any abnormalities | 4 | 1 | 1 | 0 | 5* |
| % Examined Yolk Sacs with any abnormalities | 16.0% | 8.3% | 6.7% | 0.0% | 33.3% |

\*Mipomersen-related observation.
†Although no heart observations were recorded at 100 µM in the examined embryos, this region is considered impacted by mipomersen due to the dose-response seen at lower doses and the high percentage (68.8%) of grossly malformed embryos at this dose. Heart abnormalities likely occurred in grossly malformed embryos that were not captured in the data because grossly malformed embryos were not examined.

Following this evaluation, all embryos and their extra-embryonic membranes were fixed for 24 hours in 10% neutral buffered formalin (NBF), initially orientated in HistoGel (Catalog # HG-4000-012, Epredia), processed and embedded as formalin fixed paraffin embedded (FFPE) samples for potential miRNAscope analysis.

Animal work was performed at GSK, Collegeville, Pennsylvania, USA^15^. All studies were conducted according to GSK’s Policy on the Care, Welfare and Treatment of Lab Animals and reviewed by the Institutional Animal Care and Use Committee at GSK or by the ethical review process at the institution where the work was performed. GSK is committed to the replacement, reduction, and refinement of animal studies (3Rs). Non-animal models and alternative technologies are part of our strategy and employed where possible. When animals are required, application of robust study design principles and peer review minimizes animal use, reduces harm, and improves benefit in studies.

### Statistical Analysis for WEC Data

Power calculations showed that using 16 rat or rabbit embryos will give a confidence rate of 86% for a true incidence rate of 20%. While there is no industry standard for group size in WEC studies, an N = 12 has been used in a streamlined rat WEC assay to classify teratogenic potential of pharmaceutical compounds with high predictivity^16^. For quantitative data statistical significance was determined using the Kruskal-Wallis test (2-tailed) for somite number and Dunnett’s test for yolk sac size and embryo size. Statistical analysis was not performed for qualitative morphological observations.

### miRNAscope

Detection of the unlabeled mipomersen oligonucleotide was carried out using an automated miRNAscope *in situ* hybridization assay, as previously described^17^. Formalin-fixed paraffin-embedded (FFPE) tissue sections cut at 5μm, onto positively charged slides, were stained according to the manufacturer’s recommendations on an automated staining platform (Leica BOND RX, Leica) using a mild heated target retrieval (88° C for 15 min).

Tissues were checked for global mRNA integrity using both Universal positive control (RNU6) (Cat # 727878-S1) and negative control (Scramble) (Cat # 727888-S1) probes and then with the target specific probe that detects the mipomersen antisense strand (Cat # 1763658-S1). Whole slide images were scanned at 40X (Nanozoomer S360, Hamamatsu).

## Results

### Rat WEC

Rat embryos were treated from GD 9 to 11 with vehicle (n=30), 1, 10, 50 or 100 µM mipomersen (n=16). All control embryos and embryos treated with ≤50 µM were able to be examined. Related to treatment, there were 11 (68.8%) grossly malformed and 1 (6.3%) dead embryos at 100µM mipomersen. Therefore, only 4 embryos at 100 µM mipomersen were able to be examined (Figure 1A, Table 1).

**Figure 1.**
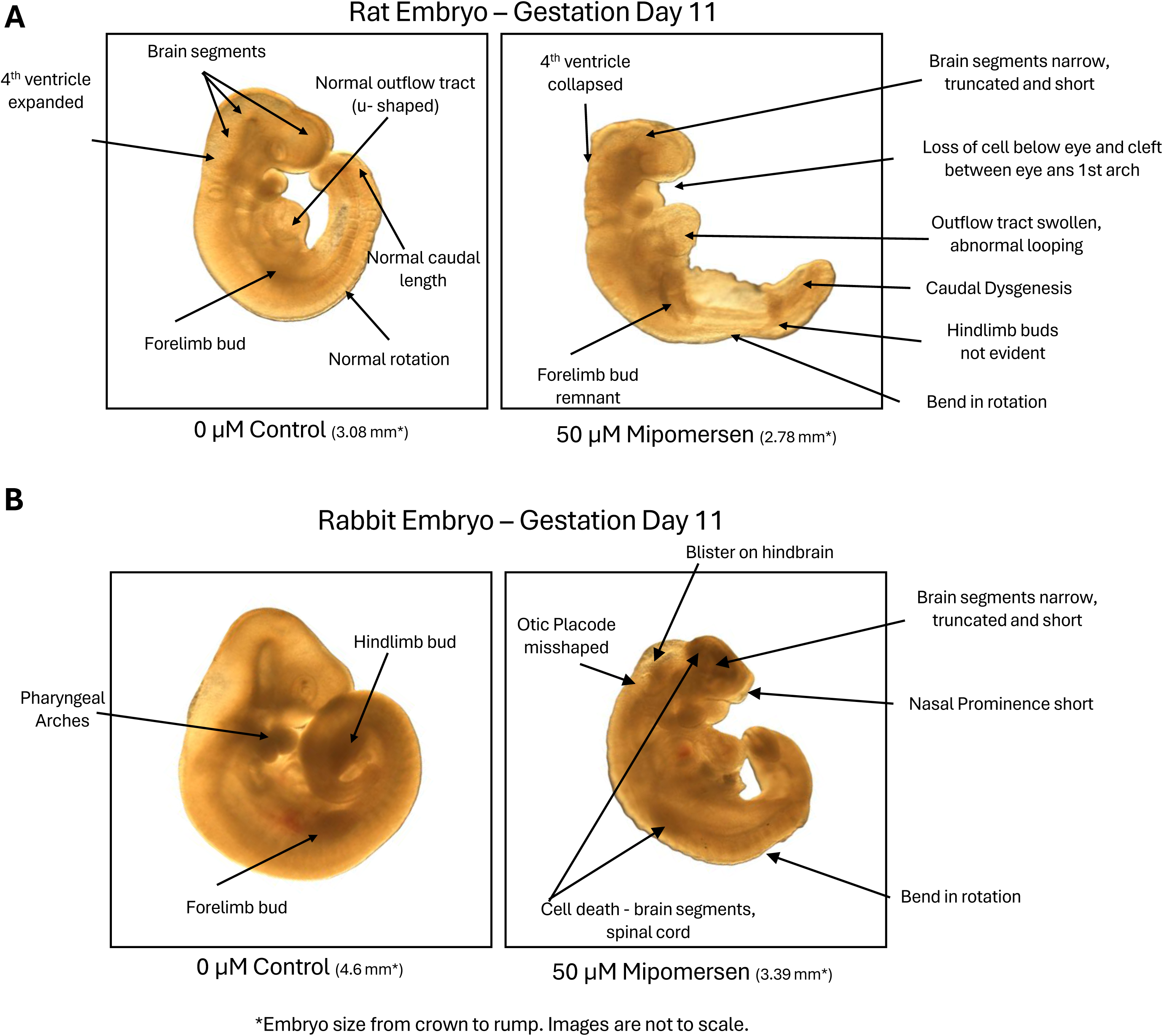

Only 1 of 4 embryos examined at 100 µM mipomersen had somites that were able to be visualized and counted. This embryo had a count of 23 somites, which is below the concurrent control mean of 27 and historical control mean of 26.5 (Supplemental Table 1). Statistical analysis was not performed on somite number due to low sample size.

Mipomersen-related effects on somites also occurred at 50 µM; the mean somite number was 25 (0.92X of control). There was mipomersen-related statistically significant, dose-dependent decrease in mean visceral yolk sac size (VYS) at 50 and 100 µM (0.93X and 0.89X of control and p = 0.005 and 0.02, respectively). There was mipomersen-related dose-dependent decrease in embryo size at 50 and 100 μM (0.93X and 0.81X of control, respectively), but statistical significance was only achieved at 100 μM (p = 0.0002) (Table 2). There were no statistically significant, mipomersen-related differences in mean VYS size, mean embryo size or somite number ≤10 μM.

**Table 2.**

| Rat WEC Quantitative Data Summary |  |  |  |  |  |  |
| --- | --- | --- | --- | --- | --- | --- |
|  |  | Dose Level |  |  |  |  |
| | | 0 μM | 1 μM | 10 μM | 50 μM | 100 μM |
| Rat Visceral Yolk Sac Size (mm) | Min. Yolk Sac Size | 3.07 | 3.08 | 2.80 | 2.71 | 2.83 |
|  | Average Yolk Sac Size | 3.53 | 3.53 | 3.44 | 3.27 | 3.14 |
|  | Max. Yolk Sac Size | 3.97 | 3.91 | 3.84 | 3.65 | 3.38 |
|  | X of Control | 1.00 | 1.00 | 0.97 | 0.93**† | 0.89**† |
| Rat Embryo Size (mm) | Min. Embryo Size | 2.51 | 2.79 | 2.60 | 2.07 | 1.78 |
|  | Average Embryo Size | 2.96 | 3.04 | 2.91 | 2.76 | 2.38 |
|  | Max. Embryo Size | 3.37 | 3.33 | 3.22 | 3.41 | 2.86 |
|  | X of Control | 1.00 | 1.03 | 0.98 | 0.93† | 0.81**† |
| Rat Somite Number | Min. Somite Number | 21 | 26 | 23 | 20 | 23 |
|  | Average Somite Number | 27 | 27 | 27 | 25 | 23 |
|  | Max. Somite Number | 28 | 28 | 28 | 27 | 23 |
|  | X of Control | 1.00 | 1.00 | 0.99 | 0.92**† | 0.86*† |
| Rabbit WEC Quantitative Data Summary |  |  |  |  |  |  |
|  |  | Dose Level |  |  |  |  |
| | | 0 μM | 1 μM | 10 μM | 50 μM | 100 μM |
| Rabbit Visceral Yolk Sac Size (mm) | Min. Yolk Sac Size | 4.39 | 3.85 | 3.81 | 4.40 | 4.19 |
|  | Average Yolk Sac Size | 5.50 | 5.14 | 5.70 | 5.72 | 5.59 |
|  | Max. Yolk Sac Size | 7.18 | 6.20 | 6.85 | 7.09 | 6.52 |
|  | X of Control | 1.00 | 0.93 | 1.04 | 1.04 | 1.02 |
| Rabbit Embryo Size (mm) | Min. Embryo Size | 2.80 | 2.72 | 2.65 | 3.39 | 3.10 |
|  | Average Embryo Size | 4.35 | 4.41 | 4.28 | 4.31 | 4.31 |
|  | Max. Embryo Size | 5.08 | 5.29 | 5.32 | 5.04 | 4.87 |
|  | X of Control | 1.00 | 1.02 | 0.98 | 0.99 | 0.99 |
| Rabbit Somite Number | Min. Somite Number | 28 | 28 | 28 | 28 | 20 |
|  | Average Somite Number | 34 | 34 | 35 | 35 | 31 |
|  | Max. Somite Number | 40 | 39 | 38 | 40 | 36 |
|  | X of Control | 1.00 | 1.00 | 1.01 | 1.03 | 0.91**† |

\*Only 1 of 4 embryos examined at 100 μM mipomersen had somites that were able to be visualized and counted; therefore, statistical analysis was not performed.
\*\*Statistically significant difference ( $p < 0.05$ ).
†Mipomersen-related observation.

Mipomersen-related dose-responsive morphological abnormalities occurred at ≥10 µM.

At 100 µM, morphological abnormalities occurred in all 4 embryos that were examined in the following regions: visceral yolk sac (abnormal vasculature, blood islands and/or paleness), caudal region (narrow or moderately short), craniofacial (cleft between eye and first pharyngeal arch, loss of cells, absent mesencephalic flexure, short nasal prominence, optic vesicle and/or otic placode abnormalities), neural tube defects (4^th^ ventricle collapsed, hemorrhage, spinal cord narrow, exencephaly, short, truncated and/or narrow brain segments), fore and hind limb buds (remnant or not evident), and embryo rotation (squirrel or squirrel with tail fused to body). Abnormalities were evident in pharyngeal arches (narrow/short) for 2 (50.0%) embryos examined. Although no heart observations were recorded at 100 µM in the examined embryos, this region is considered impacted by mipomersen due to the dose-response seen at lower doses and the high percentage (68.8%) of grossly malformed embryos at this dose. Heart abnormalities likely occurred in grossly malformed embryos that were not captured in the data because grossly malformed embryos were not examined.

At 50 μM abnormalities occurred in VYS (paleness) in 2 (12.5%) embryos, caudal (mild or moderately short, remnant and dysgenesis) in 8 (50.0%) embryos, craniofacial (cleft, loss of cells, mesencephalic flexure misshapen, nasal prominence short or not evident and optic vesicle abnormalities) in 7 (43.8%) embryos, neural tube defects (4^th^ ventricle collapsed or swollen, hemorrhage, exencephaly, short, truncated and/or narrow brain segments) in 7 (43.8%) embryos, forelimb buds (remnant or small) in 10 (62.5%) embryos, hindlimb buds (not evident) in 3 (18.8%) embryos, heart (outflow tract swollen or abnormal looping, chambers not well defined) in 4 (25.0%) embryos, embryo rotation (bend or squirrel) in 7 (43.8%) embryos and pharyngeal arches (narrow/short, remnant or not evident) in 3 (18.8%) embryos. At 10 μM abnormalities were observed in the caudal region (moderately short) in 1 (6.3%) embryo, forelimb buds (small) in 2 (12.5%) embryos, and heart (outflow tract abnormal looping, atrium swollen) in 1 (6.3%) embryo (Table 1, Supplemental Table 2).

There were no mipomersen-related effects in rat embryos at 1 µM.

### Rabbit WEC

Rabbit embryos were treated from GD 9 to 11 with vehicle (n=30) or 1, 10, 50 or 100 µM mipomersen (n=16). There were 5 (16.7%) control embryos and 3 (18.8%), 1 (6.3%), and 2 (12.5%) embryos at 1, 10 and 50 μM, respectively, found outside the visceral yolk sac which were not examined (considered to be an artifact of culture). There was 1 (6.3%) grossly malformed embryo in each group at 50 and 100 μM, which were considered unrelated to treatment due to being within historical control incidence (Supplemental Table 3). There was 1 dead embryo at 1 μM, which was not considered treatment related due to lack of dose response. These embryos were not examined; therefore, the total number of embryos examined was 25 (vehicle) and 12, 15, 13, and 15 at 1, 10, 50 and 100 μM, respectively (Figure 1B, Table 1).

There was a test article-related statistically significant decrease in mean somite number at 100 μM (0.91X of control, 31 somites at 100 μM versus 34 somites controls, p = 0.004), but no effects on visceral yolk sac size or embryo size (Table 2).

Mipomersen related morphological abnormalities occurred only at the high dose of 100 μM. Effects were observed in the visceral yolk sac (collapsed, paleness, abnormal vasculature, blood island(s), blebbing/blister) in 5 (33.3%) embryos at 100 μM compared to 4 (16.0 %) control embryos. Embryo rotation defects (bend, S-shape turning or squirrel) occurred in 7 (46.7%) embryos at 100 μM compared to 5 (20.0%) control embryos.

Hindlimb bud abnormalities (small or remnant) occurred in 6 (40.0%) embryos at 100 μM compared to 3 (12.0%) control embryos. Craniofacial defects occurred in 5 (33.3%) embryos (short nasal prominence, optic vesicle observations) compared to 3 (12.0%) controls. Mipomersen related abnormalities at 100 μM also occurred in somite morphology in 3 (20.0%) embryos (not well defined, not able to visualize, blister) and heart in 3 (20.0%) embryos (abnormal out-flow tract looping, swollen pericardial sac) compared to the vehicle control group in which none of these abnormalities were observed (Table 1, Supplemental Table 4).

Due to lack of dose response and/or incidence within concurrent or historical control, there were no mipomersen-related effects in rabbit embryos at ≤50 μM (Supplemental Table 3).

### miRNAscope

To confirm exposure, automated miRNAscope *in situ* hybridization was used to visualize the distribution of mipomersen within the embryos and on the visceral yolk sac and amnion retained following measurements and examination of embryo morphology. Although morphological examinations were not performed on grossly malformed samples, miRNAscope was performed on sections through 12 rat embryos at 100 μM mipomersen dose, 16 at 50 μM, 16 at 10 μM, 15 at 1 μM and 12 controls. Additionally, 9 rabbit embryos at 100 μM mipomersen dose, 11 each at 1, 10, and 50 μM and 13 control rabbit embryos were assessed. For all samples, an initial Haematoxylin and Eosin stain was generated and serial sections stained with the universal positive and negative control probes to quality control for overall morphology and global RNA quality (data not shown).

The embryo and the extra-embryonic membranes of rat and rabbit control embryos were all completely negative for the ONT (Figures 2A and 3A). At the 1μM dose, all rat and rabbit embryos demonstrated strong staining on the visceral yolk sac endoderm, the outermost layer of the yolk sac, but staining was noticeably absent from the embryonic mesoderm-derived endothelial, mesothelial and hematopoietic cells, and the amnion. The embryo itself was negative for the oligo signal with the notable exception of the connection between the vitelline blood vessels and the embryonic gut in both rats and rabbits (Figures 2B and 3B).

**Figure 2.**
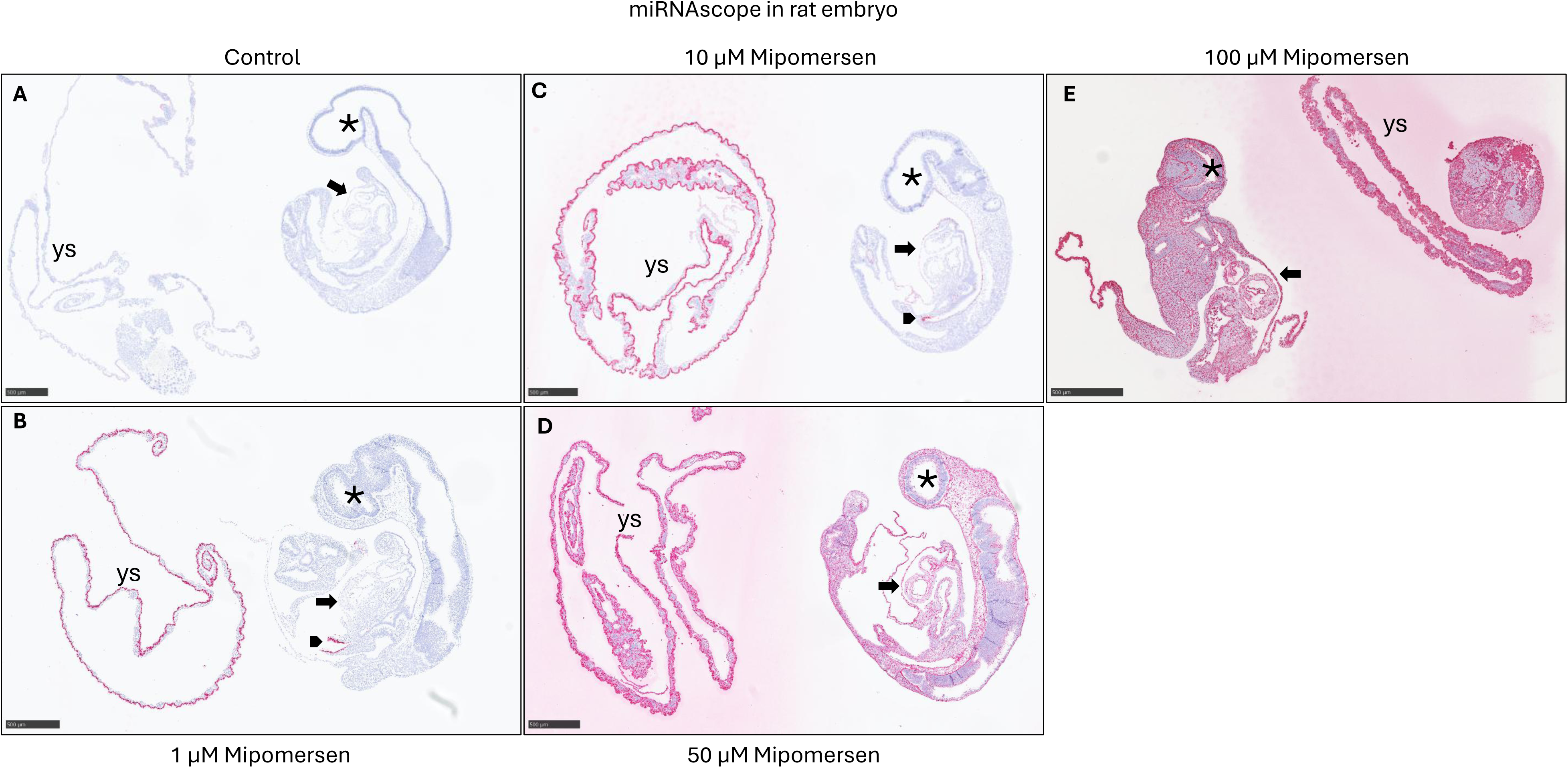
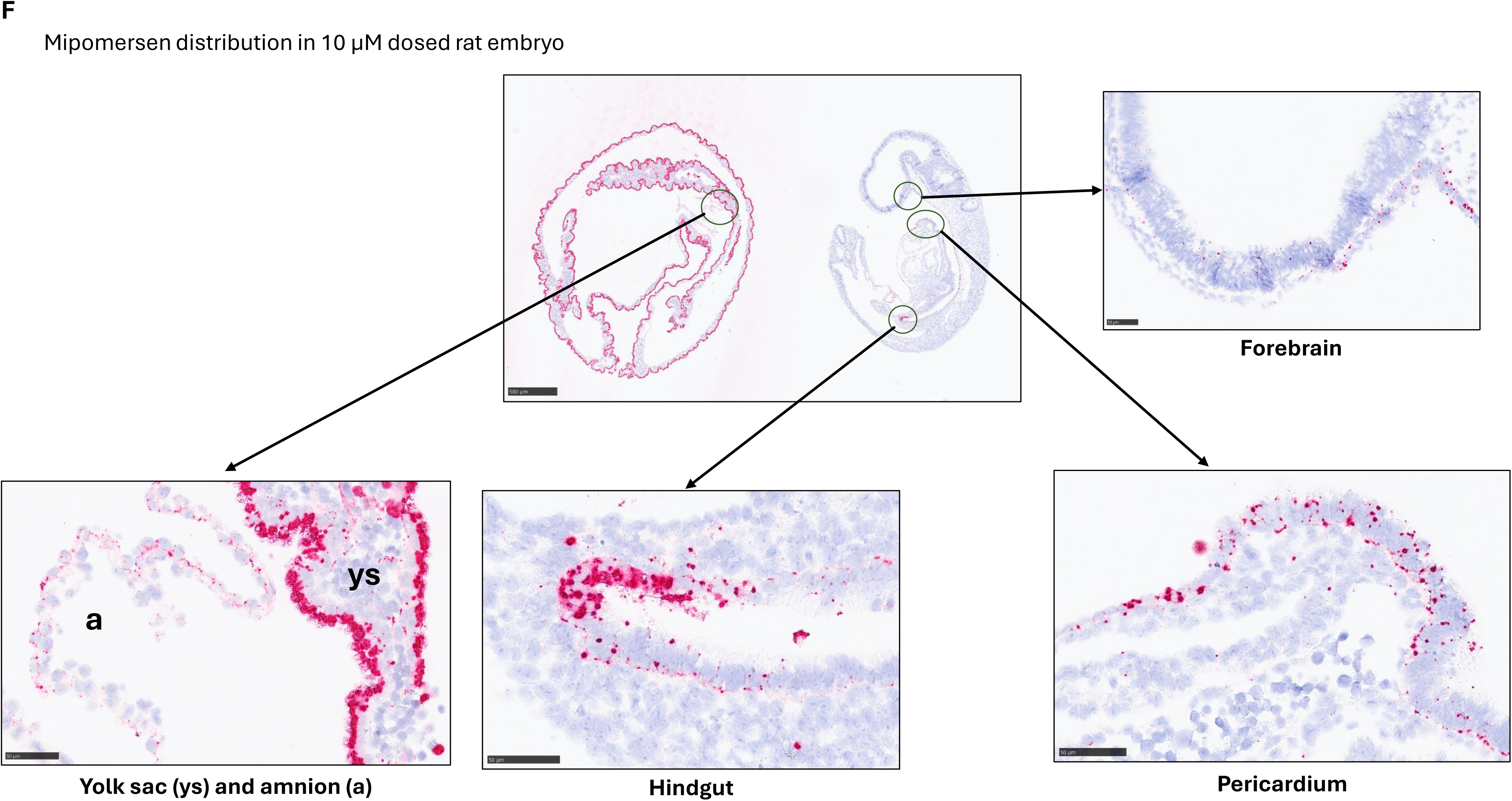
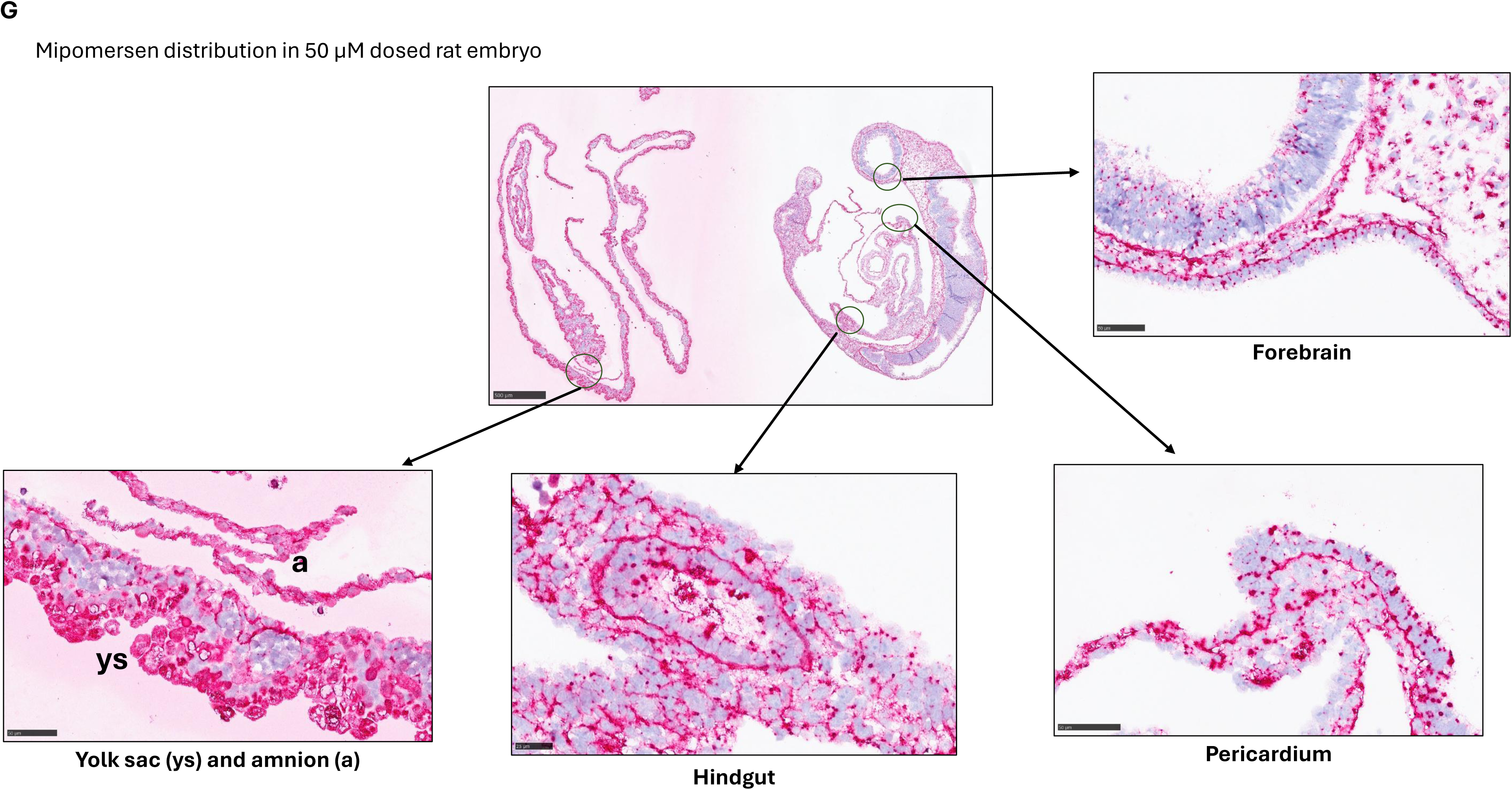

**Figure 3.**
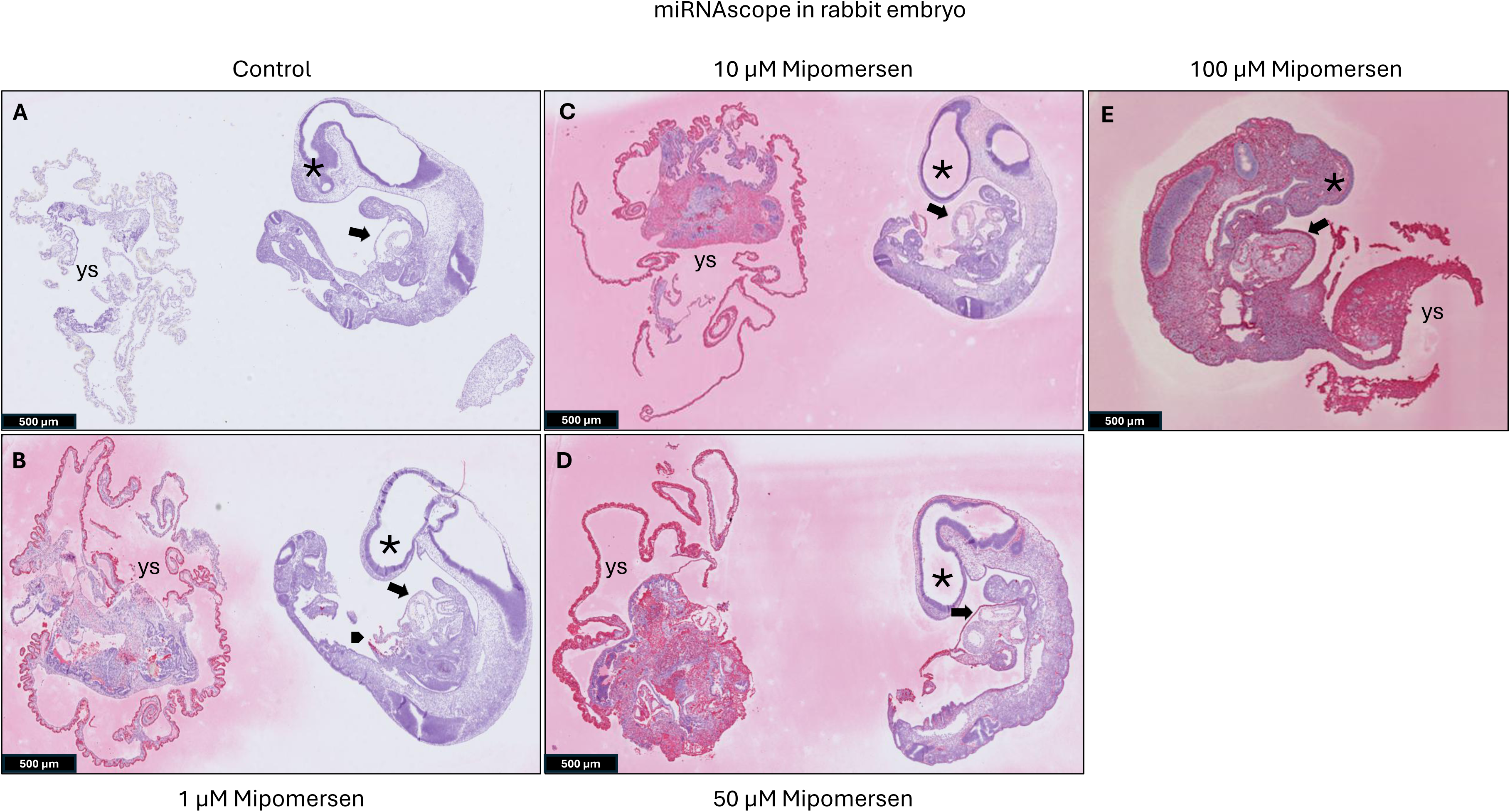
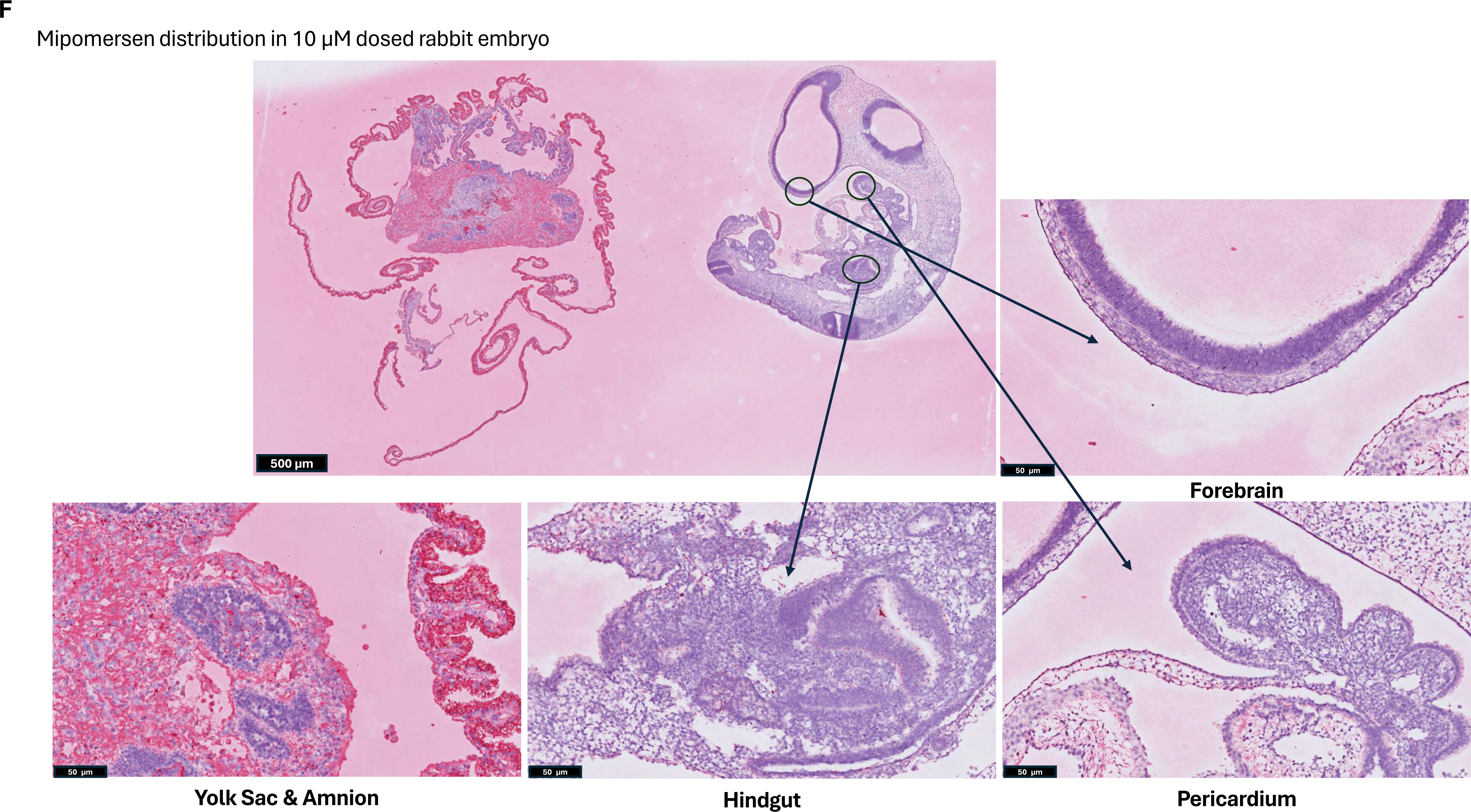
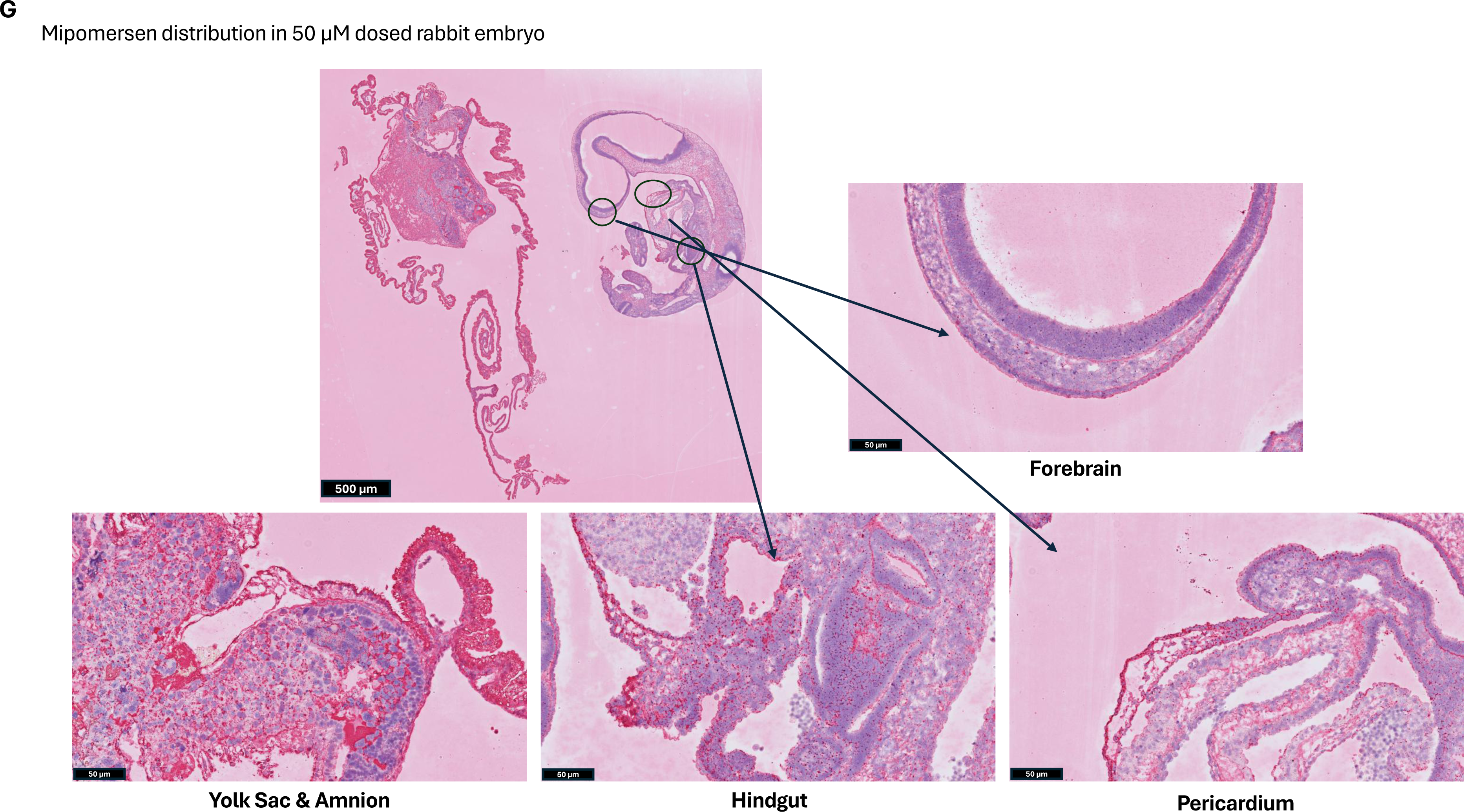

At 10 µM, the rat yolk sac staining was largely the same as at 1 µM, although weak signal was seen in the endothelial cells and in the amnion itself. However, the rabbit yolk sac staining at 10 µM showed higher mipomersen expression as compared to 1 µM treated embryos (Figures 2C and 3C). ONT was detected on the pericardium wall of the developing heart, the surface ectoderm of the craniofacial region, and at low levels in the forebrain neuroepithelium in both rat and rabbit embryos (Figures 2F and 3F). Most notable at this stage was the appearance of the ONT in the hindgut endoderm (Figures 2F and 3F).

At both the 50 µM and the 100 µM doses for rat and rabbits, all embryos and extra-embryonic membranes were heavily stained in all epithelial and mesenchymal cell layers throughout the samples (Figures 2D, E, G and 3D, E, G).

## Discussion

After culture in media containing 0, 1, 10, 50 or 100 µM mipomersen, both rat and rabbit embryos showed dose-responsive changes in yolk sac and/or embryo parameters suggestive of exposure to the ONT. At the high dose of 100 µM, rats were more severely impacted as compared to rabbits with most rat embryos being grossly malformed and those that were examinable were characterized by decreased yolk sac and embryo size, decreased somite number, and changes to yolk sac and embryo morphology. Most rabbit embryos at 100 µM were examinable and were less severely impacted, only showing a decrease in somite number and changes to embryo morphology. In rats, we observed a clear dose–response relationship. No effects were noted at the lowest dose of 1 µM even with demonstrated exposure in the vitelline blood vessels and embryonic gut, with progressively greater effects seen at each higher dose. In rabbits, a dose–response was evident only at the high dose, where test item effects were observed despite demonstrated exposure in the embryo at all dose levels.

The differences observed between species may be partially due to differences in off-target profiles. Since mipomersen is not pharmacologically active in rat or rabbit^9^, all observed effects are presumably due to off-target toxicity, which may be hybridization dependent or independent, and/or chemical/physical property-related toxicity.

Rats and rabbits also have structural and functional differences in their extraembryonic membranes^18^. Rat embryos are completely enclosed by the yolk sac during the culture period and have a Reichert’s membrane which is rodent-specific, whereas rabbit embryos are not completely enveloped by the yolk sac until after the culture period on approximately GD 12-13^13,14,18,19^. Unlike humans, rats and rabbits have an inverted yolk sac meaning that the endodermal surface, which is specialized for nutrient absorption, directly interfaces with the maternal environment *in vivo* or with the cell culture media in WEC^20,21^.

The yolk sac endoderm is highly endocytotically active and supplies nutrients via histiotrophic nutrition. Disruption of these processes at higher doses in rat WEC could have contributed to the higher incidence of abnormalities, as compared to rabbits which were not completely enveloped in yolk sac during the culture period, as interference with histiotrophic trafficking has been shown to cause malformations and resorptions *in vivo*^20^.

Automated miRNAscope *in situ* hybridization was used to confirm direct mipomersen exposure inside the embryos at all dose levels, including dose levels at which no embryo or VYS changes occurred. This suggests that ONTs must be able to get into the embryo without inherently causing toxicity. At lower doses, the allantois and heart were the first structures to show positive signal, suggesting that ONTs may enter the umbilical vasculature and pass into embryonic circulation in a dose dependent manner until saturating at higher doses where nearly ubiquitous mipomersen distribution was seen throughout all major tissues and structures in all dosed rat and rabbit embryos.

Endocytosis may allow ONTs to be taken up by the endodermal surface of the yolk sac and pass into the umbilical vasculature, as ONTs are often modified to enhance endocytosis and improve resistance to nuclease degradation, leading to longer half-lives and increased stability within the target cell^8^.

ONTs have properties of both small molecules and biologics. In the absence of internationally harmonized regulatory guidance specific to ONTs, guidelines for both small molecules (i.e., ICH M3) and biologics (i.e., ICH S6) as well as DART-specific principles [i.e., ICH S5(R3)] are typically considered during development of ONTs as potential medicines^22–24^. Additionally, the Guideline for Preclinical Safety Assessment of Oligonucleotide Therapeutics issued by the Pharmaceuticals and Medical Devices Agency (PMDA) in Japan and the draft Nonclinical Safety Assessment of Oligonucleotide-Based Therapeutics Guidance for Industry issued by the FDA should be considered when appropriate^25,26^.

When a candidate ONT is not pharmacologically active in preclinical species, the consensus is that *in vivo* DART study designs should include several dose levels of the clinical candidate in addition to one group that utilizes an appropriate species-specific surrogate molecule for hazard identification^1^. This approach was implemented for the mipomersen DART evaluation in the mouse combined fertility and early embryonic development and rabbit embryo-fetal development studies^9^. There was no evidence of impaired fertility or embryo-fetal development in mice or rabbits at doses up to 2x and 5x clinical exposure, respectively. This lack of effects as compared to our WEC results could be because mipomersen was unable to access embryonic tissue *in vivo* as it was not detectable in the fetal liver or kidney but was detectable in the mouse and rabbit placenta; alternatively, the method of detection may not have been sensitive enough and mipomersen may have been present at a low concentrations in the embryo-fetal compartment without any effects *in vivo*. When using surrogate molecules only hazard identification is possible and not risk assessment. As the current data demonstrates the ability of ONT to access embryonic tissues via WEC, it suggests that WEC assays could be used as a hazard identification assay for early embryonic development, potentially replacing the hazard identification surrogate arm of an *in vivo* study. This would reduce animal use and limit the amount of surrogate molecule quantities needed. WEC studies may provide more robust hazard identification, as compared to embryo-fetal development studies including a surrogate group, as embryonic exposure in WEC may be confirmed while maternal components may limit ONT transfer to the embryo-fetal compartment *in vivo*.

Mipomersen exposure in placental tissue in the DART studies was 23 µg/g and 15 µg/g in mice and rabbits at the high doses of 25 mg/kg/dose and 15 mg/kg/dose, respectively. The dose of 100 µM mipomersen in WEC media represents approximately 33x or 50x the placental exposure in mice and rabbits, respectively, before accounting for protein binding. Mipomersen is expected to be >85% protein-bound^27^. Since the culture media is abundant in proteins, a relatively high concentration of 100 µM mipomersen was selected to ensure that adequate levels of unbound mipomersen would be available to enter embryos. Lower doses of 1, 10 and 50 µM mipomersen were selected to evaluate a dose response and the low doses are approximately at parity with the placental exposures observed during the *in vivo* DART studies.

Because ONT properties vary, future WEC experiments should assess distribution within embryos for different antisense chemistries, as well as different tool ONT classes (e.g. small interfering RNA, microRNA mimics, transfer RNA, decoys, and aptamers). Future studies should also utilize ONTs that are pharmacologically active in the test species, enabling assessment of productive uptake, which was not possible in the current study due to lack of target engagement. Knockdown levels of the target protein should be measured to confirm functional effect of ONT treatment. Consideration should be given to selecting ONTs that target mRNAs encoding proteins with known roles in embryo-fetal development and protein expression during the culture period. These studies will further validate the utility of WEC models for hazard identification in this context.

In addition, *in vivo* studies administering ONTs should assess placental and embryo–fetal distribution using an *in situ* hybridization-type approach or immunohistochemistry to provide single-cell resolution and spatial context^28^. Most previous studies have used tissue lysate–based quantification methods such as LC–MS/MS, hybridization ELISA or Q-PCR, which measure concentrations in a diluted homogenate. These methods may or may not offer the level of sensitivity required to reliably detect ONTs in fetal samples. ONTs can enter umbilical vasculature and embryos in WEC and placental exposure has been confirmed *in vivo*. Therefore, it is plausible that ONTs are entering the embryo-fetal compartment *in vivo*, unless the placenta is acting as a barrier to prevent exposure. A thorough investigation of ONT distribution in placenta is warranted to determine if evidence of a barrier can be visualized. At early developmental stages before the placenta develops, the human yolk sac provides initial nutrient transfer and gives rise to the first blood cells of the embryo^29^. ASGR1 is expressed in human yolk sac endoderm at approximately 8 weeks post-conception^30^. ASGR1 is a receptor that is highly expressed in hepatocytes and specifically binds the GalNAc molecule to achieve targeted exposure of GalNAc siRNAs in the liver; however, ASGR1 expression in human yolk sac could lead to early embryonic exposure. This further highlights the need for additional investigations using a variety of ONT classes.

### Conclusions

We have demonstrated that WEC experiments involving mipomersen, a 2’-O-methoxyethyl phosphorothioated ASO, do not require microinjection or assisted transfection, enabling developmental hazard identification of off-target toxicity by directly exposing rat and rabbit embryos to this ONT in culture media. Hazard identification is especially important if embryo-fetal exposure may be limited *in vivo*, as it is not understood if transfer of ONTs between the maternal and embryo-fetal compartments in non-clinical species is predictive of that in humans. While WEC data cannot contribute to risk assessment due to the absence of maternal factors, identification of potential hazards may help inform prescribers and patients. The data suggest that current embryo-fetal development studies in two species for ONT development could be replaced with a hazard identification WEC study with the appropriate surrogate molecule and an in vivo second species using the clinical candidate to identify non-specific toxicity, thereby reducing animal use. The utility of WEC as a NAM for assessing developmental hazards of ONTs should be considered as the new International Council for Harmonisation of Technical Requirements for Pharmaceuticals for Human Use (ICH) S13 guidelines are drafted for the nonclinical safety evaluation of ONTs^31^.

## Acknowledgements

The authors would like to thank Heather Laurie-Williams, Joseph Hosford and Doug Fuerst (GSK Drug Substance Supply) for the synthesis and supply of mipomersen.

## Authorship confirmation/contribution statement (CReDiT format)

The following authors contributed to conceptualization and writing the original draft: Sara Bender, Dinesh, Stanislaus, Bradley Spencer-Dene. The following authors contributed to methodology, formal analysis, investigation, resources, data curation, and visualization: Joyce Rendemonti, Stanley Robinson, Sharon Chapman, Bradley Spencer-Dene, Sreenivas Nannapaneni. The following authors contributed to review and editing: Sara Bender, Joel Parry, Bradley Spencer-Dene, Dinesh Stanislaus, Joyce Rendemonti, Stanley Robinson, Sharon Chapman, Sreenivas Nannapaneni. Sara Bender was responsible for supervision and project administration.

## References

1. Hannas BR, Bender SM, Blasi E, et al. Developmental and Reproductive Toxicity Testing Strategies for Oligonucleotides: A Workshop Proceedings. Nucleic Acid Therapeutics 2025;35(3):93–113, doi:10.1089/nat.2025.0013

2. Clarke MT, Remesal L, Lentz L, et al. Prenatal delivery of a therapeutic antisense oligonucleotide achieves broad biodistribution in the brain and ameliorates Angelman syndrome phenotype in mice. Mol Ther 2024;32(4):935–951, doi:10.1016/j.ymthe.2024.02.004

3. Augustine KA, Liu ET, Sadler TW. Antisense inhibition of engrailed genes in mouse embryos reveals roles for these genes in craniofacial and neural tube development. Teratology 1995;51(5):300–10, doi:10.1002/tera.1420510506

4. Båvik C, Ward SJ, Chambon P. Developmental abnormalities in cultured mouse embryos deprived of retinoic by inhibition of yolk-sac retinol binding protein synthesis. Proc Natl Acad Sci U S A 1996;93(7):3110–4, doi:10.1073/pnas.93.7.3110

5. Foerst-Potts L, Sadler TW. Disruption of Msx-1 and Msx-2 reveals roles for these genes in craniofacial, eye, and axial development. Dev Dyn 1997;209(1):70–84, doi:10.1002/(sici)1097-0177(199705)209:1<70::Aid-aja7>3.0.Co;2-u

6. Tejedor G, Laplace-Builhé B, Luz-Crawford P, et al. Whole embryo culture, transcriptomics and RNA interference identify TBX1 and FGF11 as novel regulators of limb development in the mouse. Sci Rep 2020;10(1):3597, doi:10.1038/s41598-020-60217-w

7. Geary RS, Norris D, Yu R, et al. Pharmacokinetics, biodistribution and cell uptake of antisense oligonucleotides. Advanced Drug Delivery Reviews 2015;87(46-51, 10.1016/j.addr.2015.01.008

8. Migliorati JM, Liu S, Liu A, et al. Absorption, Distribution, Metabolism, and Excretion of US Food and Drug Administration-Approved Antisense Oligonucleotide Drugs. Drug Metab Dispos 2022;50(6):888–897, doi:10.1124/dmd.121.000417

9. Application Number 203568Orig1s000 Pharmacology Review(s). 2012.

10. New DA. Whole-embryo culture and the study of mammalian embryos during organogenesis. Biol Rev Camb Philos Soc 1978;53(1):81–122, doi:10.1111/j.1469-185x.1978.tb00993.x

11. Downs KM, Davies T. Staging of gastrulating mouse embryos by morphological landmarks in the dissecting microscope. Development 1993;118(4):1255–66, doi:10.1242/dev.118.4.1255

12. Brown NA, Fabro S. Quantitation of rat embryonic development in vitro: a morphological scoring system. Teratology 1981;24(1):65–78, doi:10.1002/tera.1420240108

13. Carney EW, Tornesi B, Keller C, et al. Refinement of a morphological scoring system for postimplantation rabbit conceptuses. Birth Defects Res B Dev Reprod Toxicol 2007;80(3):213–22, doi:10.1002/bdrb.20118

14. Pitt JA, Carney EW. Development of a morphologically-based scoring system for postimplantation New Zealand White rabbit embryos. Teratology 1999;59(2):88–101, doi:10.1002/(sici)1096-9926(199902)59:2<88::Aid-tera3>3.0.Co;2-7

15. National Research Council (US) Comitte for the Update of the Guide for the Care and Use of Laboratory Animals. Guide for the Care and Use of Laboratory Animals. 8^th^ edition. Washington (DC): National Academies Press (US); 2011. doi: 10.17226/12910

16. Zhang C, Cao J, Kenyon JR, et al. Development of a streamlined rat whole embryo culture assay for classifying teratogenic potential of pharmaceutical compounds. Toxicol Sci 2012;127(2):535–46, doi:10.1093/toxsci/kfs112

17. Spencer-Dene B, Mukherjee P, Alex A, et al. Localization of unlabeled bepirovirsen antisense oligonucleotide in murine tissues using in situ hybridization and CARS imaging. Rna 2023;29(10):1575–1590, doi:10.1261/rna.079699.123

18. Ozolinš TRS. Rabbit Whole Embryo Culture. In: Developmental Toxicology: Methods and Protocols. (Hansen JM, Winn LM. eds.) Springer New York: New York, NY; 2019; pp. 219-233.

19. Furukawa S, Tsuji N, Sugiyama A. Morphology and physiology of rat placenta for toxicological evaluation. J Toxicol Pathol 2019;32(1):1–17, doi:10.1293/tox.2018-0042

20. Holson JF, Stump DG, Pearce LB, et al. Mode of action: yolk sac poisoning and impeded histiotrophic nutrition--HBOC-related congenital malformations. Crit Rev Toxicol 2005;35(8-9):739–45, doi:10.1080/10408440591007412

21. Stokes PA, Vardy PH, Mc BW. Advances in Rabbit Embryo Culture during Organogenesis. Dev Growth Differ 1981;23(6):623–627, doi:10.1111/j.1440-169X.1981.00623.x

22. ICH Harmonised Guideline M3(R2) Guidance on Nonclinical Safety Studies for the Conduct of Human Clinical Trials and Marketing Authorization for Pharmaceuticals 2009.

23. ICH Harmonised Guideline S6(R1) Preclinical Safety Evaluation of Biotechnology-Derived Pharmaceuticals 2011.

24. ICH Harmonised Guideline S5(R3) Detection of Reproductive and Developmental Toxicity for Human Pharmaceuticals 2020.

25. PMDA. Guideline for preclinical safety assessment of oligonucleotide therapeutics. 2020.

26. U.S. Department of Health and Human Services FaDA, Center for Drug Evaluation and Research (CDER). Nonclinical Safety Assessment of Oligonucleotide-Based Therapeutics Guidance for Industry - Draft Guidance 2024.

27. Jiang R, Hooshfar S, Eno MR, et al. Factors Influencing ADME Properties of Therapeutic Antisense Oligonucleotides: Physicochemical Characteristics and Beyond. Current Drug Metabolism 2023;24(7):536–552, 10.2174/1389200224666230418092626

28. Fial I, Farrier SA, Chimento DP, et al. Characterizing Antibodies Targeting Antisense Oligonucleotide Phosphorothioate and 2’-O-Methoxyethyl Modifications for Intracellular Trafficking and Biodistribution Studies. Nucleic Acid Ther 2025;35(4):168–181, doi:10.1177/21593337251361396

29. Ross C, Boroviak TE. Origin and function of the yolk sac in primate embryogenesis. Nature Communications 2020;11(1):3760, doi:10.1038/s41467-020-17575-w

30. Goh I, Botting RA, Rose A, et al. Yolk sac cell atlas reveals multiorgan functions during human early development. Science 2023;381(6659):eadd7564, doi:doi:10.1126/science.add7564

31. S13 EWG: Non-clinical Safety Evaluation of Oligonucleotide-based Therapeutics. Final Concept Paper 2024. https://database.ich.org/sites/default/files/ICH_S13EWG_Concept_Paper_2024_1028.pdf

