## Supplemental Tables for "Rat and rabbit whole embryo culture as a new approach method for unlabeled therapeutic antisense oligonucleotide hazard identification with no requirement for microinjection or assisted transfection"

### Supplemental Table 1.

| Rat WEC Quantitative Data – Historical Control Data* | | | | | | | | | | | |
|---|---|---|---|---|---|---|---|---|---|---|---|
| Visceral Yolk-Sac Sizes (mm)* | | | | Embryo Sizes (mm)* | | | | Number of Somites* | | | |
| WEC Study | Min. Yolk-Sac Size | Average Yolk-Sac Size | Max. Yolk-Sac Size | WEC Study | Min. Embryo Size | Average Embryo Size | Max. Embryo Size | WEC Study | Min. Somite Number | Average Somite Number | Max. Somite Number |
| WEC 1 | 3.69 | 3.78 | 3.88 | WEC 1 | 2.97 | 3.17 | 3.39 | WEC 1 | 26 | 26.3 | 27 |
| WEC 2 | 3.37 | 3.67 | 4.08 | WEC 2 | 2.91 | 3.10 | 3.48 | WEC 2 | 25 | 26.5 | 28 |
| WEC 3 | 3.51 | 3.75 | 4.08 | WEC 3 | 2.86 | 3.10 | 3.32 | WEC 3 | 24 | 26.2 | 27 |
| WEC 4 | 3.43 | 3.80 | 4.16 | WEC 4 | 2.87 | 3.17 | 3.52 | WEC 4 | 26 | 26.5 | 28 |
| WEC 5 | 3.39 | 3.73 | 4.09 | WEC 5 | 2.91 | 3.09 | 3.35 | WEC 5 | 25 | 26.3 | 28 |
| WEC 6 | 3.56 | 3.70 | 3.89 | WEC 6 | 3.04 | 3.13 | 3.19 | WEC 6 | 26 | 26.5 | 27 |
| WEC 7 | 3.23 | 3.68 | 4.11 | WEC 7 | 2.83 | 3.14 | 3.32 | WEC 7 | 25 | 26.3 | 27 |
| WEC 8 | 3.22 | 3.73 | 4.22 | WEC 8 | 2.92 | 3.19 | 3.37 | WEC 8 | 26 | 26.6 | 27 |
| WEC 9 | 3.40 | 3.57 | 3.77 | WEC 9 | 2.88 | 3.08 | 3.34 | WEC 9 | 25 | 26.6 | 28 |
| WEC 10 | 3.46 | 3.77 | 4.03 | WEC 10 | 2.90 | 3.18 | 3.42 | WEC 10 | 26 | 27.1 | 28 |
| WEC 11 | 3.56 | 3.82 | 4.13 | WEC 11 | 2.91 | 3.19 | 3.42 | WEC 11 | 26 | 27.2 | 28 |
| WEC 12 | 3.19 | 3.70 | 4.18 | WEC 12 | 2.43 | 3.15 | 3.55 | WEC 12 | 25 | 26.8 | 28 |
| WEC 13 | 3.11 | 3.50 | 3.89 | WEC 13 | 2.87 | 3.05 | 3.34 | WEC 13 | 25 | 26.1 | 27 |
| WEC 14 | 3.28 | 3.76 | 4.16 | WEC 14 | 2.62 | 3.16 | 3.50 | WEC 14 | 24 | 26.2 | 27 |
| WEC 15 | 3.52 | 3.77 | 3.99 | WEC 15 | 2.89 | 3.21 | 3.47 | WEC 15 | 25 | 26.6 | 28 |
| WEC 16 | 3.23 | 3.76 | 3.99 | WEC 16 | 2.94 | 3.18 | 3.37 | WEC 16 | 26 | 26.6 | 28 |
| WEC 17 | 3.47 | 3.77 | 4.16 | WEC 17 | 2.88 | 3.21 | 3.41 | WEC 17 | 25 | 26.6 | 28 |
| WEC 18 | 3.63 | 3.79 | 3.94 | WEC 18 | 3.02 | 3.10 | 3.17 | WEC 18 | 26 | 26.0 | 26 |
| WEC 19 | 3.36 | 3.75 | 4.22 | WEC 19 | 2.91 | 3.17 | 3.45 | WEC 19 | 25 | 26.7 | 28 |
| WEC 20 | 3.29 | 3.73 | 4.10 | WEC 20 | 2.81 | 3.17 | 3.39 | WEC 20 | 25 | 26.8 | 27 |
| WEC 21 | 3.18 | 3.69 | 4.10 | WEC 21 | 2.78 | 3.18 | 3.68 | WEC 21 | 25 | 26.8 | 28 |
| WEC 22 | 3.37 | 3.58 | 3.75 | WEC 22 | 2.81 | 3.14 | 3.32 | WEC 22 | 25 | 26.2 | 27 |
| WEC 23 | 3.51 | 3.81 | 4.14 | WEC 23 | 3.09 | 3.29 | 3.60 | WEC 23 | 26 | 27.0 | 28 |
| WEC 24 | 3.45 | 3.78 | 4.28 | WEC 24 | 2.98 | 3.19 | 3.39 | WEC 24 | 25 | 26.4 | 27 |
| WEC 25 | 3.15 | 3.55 | 3.95 | WEC 25 | 2.67 | 3.03 | 3.23 | WEC 25 | 25 | 26.1 | 27 |
| WEC 26 | 3.13 | 3.61 | 3.96 | WEC 26 | 2.75 | 3.07 | 3.37 | WEC 26 | 25 | 26.6 | 28 |
| WEC 27 | 3.45 | 3.74 | 3.99 | WEC 27 | 2.99 | 3.21 | 3.39 | WEC 27 | 25 | 26.6 | 28 |
| WEC 28 | 3.35 | 3.72 | 4.07 | WEC 28 | 2.87 | 3.13 | 3.25 | WEC 28 | 26 | 26.4 | 27 |
| WEC 29 | 3.20 | 3.56 | 3.89 | WEC 29 | 2.88 | 3.11 | 3.31 | WEC 29 | 25 | 26.4 | 27 |
| WEC 30 | 3.29 | 3.77 | 4.20 | WEC 30 | 2.74 | 3.22 | 3.56 | WEC 30 | 25 | 26.9 | 28 |
| WEC 31 | 3.26 | 3.60 | 3.87 | WEC 31 | 2.74 | 3.05 | 3.30 | WEC 31 | 25 | 26.3 | 27 |
| WEC 32 | 3.18 | 3.75 | 4.02 | WEC 32 | 2.88 | 3.15 | 3.33 | WEC 32 | 26 | 26.6 | 28 |
| **Overall Range** | **3.11** | **NA** | **4.28** | **Overall Range** | **2.43** | **NA** | **3.68** | **Overall Range** | **24** | **NA** | **28** |
| **Overall Mean**** | **NA** | **3.71** | **NA** | **Overall Mean**** | **NA** | **3.15** | **NA** | **Overall Mean**** | **NA** | **26.5** | **NA** |

*Data includes studies from the past 5 years.

**Overall mean is the mean of all embryos in the historical control data.

NA = not applicable

### Supplemental Table 2.

| Rat WEC Morphology Detailed Observations | | | | | |
|---|---|---|---|---|---|
| **Dose Level** | **0 µM** | **1 µM** | **10 µM** | **50 µM** | **100 µM** |
| **Able to Examine** | | | | | |
| Yes | 30 | 16 | 16 | 16 | 4 |
| No | 0 | 0 | 0 | 0 | 12 |
| Dead | 0 | 0 | 0 | 0 | 1 |
| Grossly Malformed | 0 | 0 | 0 | 0 | 11 |
| Not in bottle | 0 | 0 | 0 | 0 | 0 |
| Embryo Outside of Yolk Sac | 0 | 0 | 0 | 0 | 0 |
| **Caudal** | | | | | |
| Not Remarkable | 30 | 16 | 15 | 8 | 0 |
| Caudal Dysgenesis | 0 | 0 | 0 | 2 | 0 |
| Other | 0 | 0 | 0 | 0 | 0 |
| Tail - Narrow | 0 | 0 | 0 | 0 | 2 |
| Tail - Remnant | 0 | 0 | 0 | 1 | 0 |
| Tail - Short; mild | 0 | 0 | 0 | 3 | 0 |
| Tail - Short; moderate | 0 | 0 | 1 | 3 | 2 |
| Total Embryos with Caudal Abnormalities | 0 | 0 | 1 | 8 | 4 |
| **Craniofacial** | | | | | |
| Not Remarkable | 30 | 16 | 16 | 9 | 0 |
| Blister | 0 | 0 | 0 | 0 | 0 |
| Cleft Between Eye and First Arch | 0 | 0 | 0 | 6 | 3 |
| Lens - Not Evident | 0 | 0 | 0 | 0 | 0 |
| Loss Cells Below Eye or Between Eye and Arch | 0 | 0 | 0 | 1 | 2 |
| Mesencephalic Flexure - Misshapen/Malformed | 0 | 0 | 0 | 1 | 0 |
| Mesencephalic Flexure - Not Evident | 0 | 0 | 0 | 0 | 3 |
| Mesencephalic Flexure - Small/Narrow | 0 | 0 | 0 | 0 | 0 |
| Nasal Prominence - Not Evident | 0 | 0 | 0 | 1 | 0 |
| Nasal Prominence - Short; Moderate | 0 | 0 | 0 | 1 | 1 |
| Nasal Prom. - Short; Severe | 0 | 0 | 0 | 0 | 3 |
| Optic Vesicle - Loss of Definition | 0 | 0 | 0 | 1 | 0 |
| Optic Vesicle - Misshapen | 0 | 0 | 0 | 2 | 1 |
| Optic Vesicle - Not Evident | 0 | 0 | 0 | 1 | 1 |
| Optic Vesicle - Remnant | 0 | 0 | 0 | 0 | 0 |
| Optic Vesicle - Small | 0 | 0 | 0 | 1 | 1 |
| Other | 0 | 0 | 0 | 0 | 0 |
| Otic Placode - Displaced | 0 | 0 | 0 | 0 | 0 |
| Otic Placode - Loss of Definition | 0 | 0 | 0 | 0 | 0 |
| Otic Placode - Misshapen | 0 | 0 | 0 | 0 | 1 |

| | | | | | |
|---|---|---|---|---|---|
| Otic Placode - Not Evident | 0 | 0 | 0 | 0 | 0 |
| Otic Placode - Small | 0 | 0 | 0 | 0 | 1 |
| Optic Vesicle - Round without lens | 0 | 0 | 0 | 0 | 0 |
| Otic Placode - Round | 0 | 0 | 0 | 0 | 0 |
| Total Embryos with Craniofacial Abnormalities | 0 | 0 | 0 | 7 | 4 |
| **Forelimb Bud** | | | | | |
| Not Remarkable | 30 | 16 | 14 | 6 | 0 |
| Not Evident | 0 | 0 | 0 | 0 | 0 |
| Other | 0 | 0 | 0 | 0 | 0 |
| Remnant | 0 | 0 | 0 | 6 | 4 |
| Small | 0 | 0 | 2 | 4 | 0 |
| Total Embryos with Forelimb Bud Abnormalities | 0 | 0 | 2 | 10 | 4 |
| **Hindlimb Buds** | | | | | |
| Not Remarkable | 30 | 16 | 16 | 13 | 0 |
| Not Evident | 0 | 0 | 0 | 3 | 4 |
| Other | 0 | 0 | 0 | 0 | 0 |
| Total Embryos with Hindlimb Bud Abnormalities | 0 | 0 | 0 | 3 | 4 |
| **Heart** | | | | | |
| Not Remarkable | 27 | 16 | 14 | 12 | 4 |
| Atrium - Small/Compressed | 0 | 0 | 0 | 0 | 0 |
| Atrium - Swollen | 0 | 0 | 1 | 0 | 0 |
| Cardiac Chambers - Cell Death | 0 | 0 | 0 | 0 | 0 |
| Chambers Not Well Defined | 0 | 0 | 0 | 1 | 0 |
| Damaged | 0 | 0 | 0 | 0 | 0 |
| Dextrocardia* | 2 | 0 | 1 | 2 | 0 |
| No Heartbeat | 0 | 0 | 0 | 0 | 0 |
| Outflow Tract - Abnormal Looping | 0 | 0 | 1 | 2 | 0 |
| Outflow Tract - Kinked | 0 | 0 | 0 | 0 | 0 |
| Outflow Tract - Narrow | 0 | 0 | 0 | 0 | 0 |
| Outflow Tract - Not Evident | 0 | 0 | 0 | 0 | 0 |
| Outflow Tract - Not Well Defined | 0 | 0 | 0 | 0 | 0 |
| Outflow Tract - Swollen | 0 | 0 | 0 | 3 | 0 |
| Other | 0 | 0 | 0 | 0 | 0 |
| Pericardial Sac - Blebbing | 0 | 0 | 0 | 0 | 0 |
| Pericardial Sac - Cell Death | 0 | 0 | 0 | 0 | 0 |
| Pericardial Sac - Filled with Blood | 0 | 0 | 0 | 0 | 0 |
| Pericardial Sac – Not Evident | 0 | 0 | 0 | 0 | 0 |
| Pericardial Sac - Swollen | 1 | 0 | 0 | 0 | 0 |
| Tube-shaped | 0 | 0 | 0 | 0 | 0 |
| Ventricle - Small/Compressed | 0 | 0 | 0 | 0 | 0 |
| Ventricle - Swollen | 0 | 0 | 0 | 0 | 0 |

| | | | | | |
|---|---|---|---|---|---|
| Total Embryos with Heart Abnormalities | 3 | 0 | 2 | 4 | 0 |
| **Neural Tube** | | | | | |
| Not Remarkable | 30 | 16 | 16 | 9 | 0 |
| 4th Ventricle - Collapsed | 0 | 0 | 0 | 4 | 2 |
| 4th Ventricle - Swollen | 0 | 0 | 0 | 1 | 0 |
| Blister | 0 | 0 | 0 | 0 | 0 |
| Brain - Hemorrhage | 0 | 0 | 0 | 3 | 1 |
| Exencephaly | 0 | 0 | 0 | 3 | 1 |
| Forebrain - Cleft | 0 | 0 | 0 | 0 | 0 |
| Forebrain - Kinked | 0 | 0 | 0 | 0 | 0 |
| Forebrain - Misshaped | 0 | 0 | 0 | 0 | 0 |
| Forebrain - Narrow | 0 | 0 | 0 | 0 | 0 |
| Forebrain - Not Evident | 0 | 0 | 0 | 0 | 0 |
| Forebrain - Short | 0 | 0 | 0 | 1 | 0 |
| Forebrain - Truncated | 0 | 0 | 0 | 0 | 0 |
| Forebrain /Midbrain - Loss of Definition | 0 | 0 | 0 | 0 | 0 |
| Forebrain /Midbrain /Hindbrain - Loss of Definition | 0 | 0 | 0 | 0 | 0 |
| Forebrain /Midbrain /Hindbrain - Narrow | 0 | 0 | 0 | 0 | 0 |
| Forebrain /Midbrain /Hindbrain - Narrow, Short, Truncated | 0 | 0 | 0 | 2 | 3 |
| Forebrain /Midbrain /Hindbrain - Short | 0 | 0 | 0 | 0 | 0 |
| Forebrain /Midbrain /Hindbrain - Truncated | 0 | 0 | 0 | 0 | 0 |
| Hindbrain - Kinked | 0 | 0 | 0 | 0 | 0 |
| Hindbrain - Narrow | 0 | 0 | 0 | 0 | 0 |
| Hindbrain - Short | 0 | 0 | 0 | 1 | 0 |
| Hindbrain - Truncated | 0 | 0 | 0 | 0 | 0 |
| Midbrain - Indentation | 0 | 0 | 0 | 0 | 0 |
| Midbrain - Kinked | 0 | 0 | 0 | 0 | 0 |
| Midbrain - Narrow | 0 | 0 | 0 | 0 | 0 |
| Midbrain - Not Evident | 0 | 0 | 0 | 0 | 0 |
| Midbrain - Short | 0 | 0 | 0 | 0 | 0 |
| Midbrain - Truncated | 0 | 0 | 0 | 0 | 0 |
| Midbrain /Hindbrain - Loss of Definition | 0 | 0 | 0 | 0 | 0 |
| Open Neuropore - Caudal | 0 | 0 | 0 | 0 | 0 |
| Open Neuropore - Forebrain | 0 | 0 | 0 | 0 | 0 |
| Open Neuropore - Hindbrain | 0 | 0 | 0 | 0 | 0 |
| Other | 0 | 0 | 0 | 0 | 0 |
| Spinal Cord - Kinking | 0 | 0 | 0 | 0 | 0 |
| Spinal Cord - Lack of Neural Fold Fusion | 0 | 0 | 0 | 0 | 0 |
| Spinal Cord - Narrow | 0 | 0 | 0 | 0 | 1 |
| Spinal Cord - Swelling | 0 | 0 | 0 | 0 | 0 |
| Cell Death | 0 | 0 | 0 | 0 | 0 |

| | | | | | |
|---|---|---|---|---|---|
| Total Embryos with Neural Tube Abnormalities | 0 | 0 | 0 | 7 | 4 |
| **Pharyngeal Arches** | | | | | |
| Not Remarkable | 30 | 16 | 16 | 13 | 2 |
| 1st Arch - Cell Death | 0 | 0 | 0 | 0 | 0 |
| 1st Arch - Hemorrhage | 0 | 0 | 0 | 0 | 0 |
| 1st Arch - Narrow/Short | 0 | 0 | 0 | 1 | 0 |
| 1st Arch – Not Evident | 0 | 0 | 0 | 0 | 0 |
| 1st Arch - Remnant | 0 | 0 | 0 | 0 | 0 |
| 1st Arch - Swollen | 0 | 0 | 0 | 0 | 0 |
| 2nd Arch - Cell Death | 0 | 0 | 0 | 0 | 0 |
| 2nd Arch - Hemorrhage | 0 | 0 | 0 | 0 | 0 |
| 2nd Arch - Narrow/Short | 0 | 0 | 0 | 1 | 2 |
| 2nd Arch – Not Evident | 0 | 0 | 0 | 1 | 0 |
| 2nd Arch - Remnant | 0 | 0 | 0 | 1 | 0 |
| Fused - 1st and 2nd | 0 | 0 | 0 | 0 | 0 |
| Fused - 1st at midline | 0 | 0 | 0 | 0 | 0 |
| Other | 0 | 0 | 0 | 0 | 0 |
| Ramus - Blister | 0 | 0 | 0 | 0 | 0 |
| Ramus - Cell Death | 0 | 0 | 0 | 0 | 0 |
| Total Embryos with Pharyngeal Arch Abnormalities | 0 | 0 | 0 | 3 | 2 |
| **Rotation** | | | | | |
| Not Remarkable | 29 | 16 | 16 | 9 | 0 |
| Bend | 0 | 0 | 0 | 4 | 0 |
| Other | 0 | 0 | 0 | 0 | 0 |
| Squirrel | 0 | 0 | 0 | 3 | 3 |
| Squirrel - Tail Fused to Body | 0 | 0 | 0 | 0 | 1 |
| S-Shaped | 1 | 0 | 0 | 0 | 0 |
| Total Embryos with Rotation Abnormalities | 1 | 0 | 0 | 7 | 4 |
| **Somite Morphology** | | | | | |
| NR | 30 | 16 | 16 | 15 | 1 |
| Cobblestone | 0 | 0 | 0 | 0 | 0 |
| Compressed | 0 | 0 | 0 | 0 | 0 |
| Fused | 0 | 0 | 0 | 0 | 0 |
| Missing | 0 | 0 | 0 | 0 | 0 |
| Narrow | 0 | 0 | 0 | 0 | 0 |
| Not Able to Visualize | 0 | 0 | 0 | 0 | 3 |
| Not well Defined | 0 | 0 | 0 | 1 | 0 |
| Other | 0 | 0 | 0 | 0 | 0 |
| Rounded | 0 | 0 | 0 | 0 | 0 |
| Serrated | 0 | 0 | 0 | 0 | 0 |
| Small | 0 | 0 | 0 | 0 | 0 |

| | | | | | |
|---|---|---|---|---|---|
| Split | 0 | 0 | 0 | 0 | 0 |
| Wedge Shaped | 0 | 0 | 0 | 0 | 0 |
| Total Embryos with Somite Abnormalities | 0 | 0 | 0 | 1 | 3 |
| **Yolk Sac Morphology** | | | | | |
| Not Remarkable | 30 | 16 | 16 | 14 | 0 |
| Abnormal Vasculature | 0 | 0 | 0 | 0 | 3 |
| Blebbing/Blister | 0 | 0 | 0 | 0 | 0 |
| Blood Cells - Not Evident | 0 | 0 | 0 | 0 | 0 |
| Blood Cells - Sparse | 0 | 0 | 0 | 0 | 0 |
| Blood Island(s) | 0 | 0 | 0 | 0 | 1 |
| Collapsed | 0 | 0 | 0 | 0 | 0 |
| Discolored | 0 | 0 | 0 | 0 | 0 |
| Ectoplacental Cone - Detached | 0 | 0 | 0 | 0 | 0 |
| Ectoplacental Cone - Small | 0 | 0 | 0 | 0 | 0 |
| Invaginated | 0 | 0 | 0 | 0 | 0 |
| Necrotic | 0 | 0 | 0 | 0 | 0 |
| Not Examined - Hole in Visceral Yolk Sac | 0 | 0 | 0 | 0 | 0 |
| Other | 0 | 0 | 0 | 0 | 0 |
| Pale | 0 | 0 | 0 | 2 | 2 |
| Rough Surface | 0 | 0 | 0 | 0 | 0 |
| Total Yolk Sacs with any abnormalities | 0 | 0 | 0 | 2 | 4 |

*Dextrocardia observations were recorded in the raw data, but not summarized, because dextrocardia is considered a culture-related artifact and not a test item effect.

### Supplemental Table 3.

| Rabbit WEC Morphology – Historical Control Data* | | | | | | | | | | | | |
|---|---|---|---|---|---|---|---|---|---|---|---|---|
| **Observations** | **WEC Study** | | | | | | | | | | | |
| **Summary of Numbers** | **WEC 1** | **WEC 2** | **WEC 3** | **WEC 4** | **WEC 5** | **WEC 6** | **WEC 7** | **WEC 8** | **WEC 9** | **WEC 10** | **WEC 11** | **Grand Total** |
| No | 1 | 5 | 1 | | | 2 | 2 | 1 | 1 | 2 | | 15 |
| Yes | 15 | 26 | 9 | 12 | 10 | 8 | 12 | 9 | 15 | 22 | 16 | 154 |
| Grossly Malformed | | 2 | | | | 1 | | | | | | 3 |
| Dead | | | | | | | | | | | | 0 |
| **Caudal** | | | | | | | | | | | | |
| Not Remarkable | 13 | 26 | 9 | 12 | 8 | 8 | 12 | 8 | 15 | 22 | 15 | 148 |
| Tail - Short; mild | 1 | | | | 2 | | | 1 | | | 1 | 5 |
| Tail - Short; moderate | 1 | | | | | | | | | | | 1 |
| **Craniofacial** | | | | | | | | | | | | |
| Not Remarkable | 14 | 25 | 9 | 11 | 10 | 8 | 12 | 9 | 14 | 20 | 15 | 147 |
| Lens - Not Evident | 1 | 1 | | | | | | | 1 | | 1 | 4 |
| Mesencephalic Flexure – Not Evident | | | | | | | | | | 1 | | 1 |
| Maxillary Process shorter than 1st arch | | 1 | | | | | | | | | | 1 |
| Nasal Prominence. – Not Evident | | | | | | | | | | 1 | | 1 |
| Nasal Prominence - Short; Moderate | | | | | | | | | 1 | 1 | | 2 |
| Nasal Prominence - Short; Severe | | 1 | | | | | | | | | 1 | 2 |
| Olfactory - small indentations | | 1 | | | | | | | | | | 1 |
| Optic Vesicle - Not Evident | | | | | | | | | | 1 | | 1 |
| Optic Vesicle - Round without lens | | | | 1 | | | | | | | | 1 |
| Optic Vesicle - Small | | | | | | | | | | | 1 | 1 |
| Otic Placode - Misshapen | | | | | | | | | | | 1 | 1 |
| **Forelimb Bud** | | | | | | | | | | | | |
| Not Remarkable | 12 | 19 | 9 | 10 | 9 | 8 | 12 | 9 | 13 | 19 | 13 | 133 |
| Not Evident | | | | | | | | | 2 | | 1 | 3 |
| Remnant | 1 | 3 | | | 1 | | | | | | | 5 |
| Small | 2 | 6 | | 2 | | | | | 1 | 3 | 2 | 16 |
| **Heart** | | | | | | | | | | | | |
| Not Remarkable | 15 | 26 | 9 | 12 | 9 | 8 | 12 | 7 | 14 | 21 | 15 | 148 |
| Atrium - Small/Compressed | | | | | 1 | | | | | | | 1 |

| | | | | | | | | | | | | |
|---|---|---|---|---|---|---|---|---|---|---|---|---|
| Dextrocardia | | | | | | | | 1 | | | | 1 |
| Outflow Tract - Abnormal Looping | | | | | 1 | | | 1 | | | 1 | 3 |
| Outflow Tract - Swollen | | | | | | | | | 1 | | | 1 |
| Pericardial Sac - Swollen | | | | | | | | | | 1 | | 1 |
| Ventricle - Small/Compressed | | | | | 1 | | | | | | | 1 |
| **Hindlimb Bud** | | | | | | | | | | | | |
| Not Remarkable | 15 | 24 | 9 | 12 | 9 | 8 | 12 | 8 | 13 | 22 | 15 | 147 |
| Not Evident | | | | | 1 | | | | 1 | | | 2 |
| Remnant | | | | | | | | | 1 | | | 1 |
| Small | | 2 | | | | | | 1 | | | 1 | 4 |
| **Neural Tube** | | | | | | | | | | | | |
| Not Remarkable | 14 | 25 | 8 | 12 | 10 | 8 | 12 | 9 | 14 | 20 | 14 | 146 |
| 4th Vent. - Collapsed | 1 | | 1 | | | | | | | | 1 | 3 |
| 4th Vent. - Swollen | | 1 | | | | | | | | | 1 | 2 |
| Blister | | | | | | | | | 1 | | | 1 |
| Cell death | | | | | | | | | | | 1 | 1 |
| Forebrain - Not Evident | | | | | | | | | | 1 | | 1 |
| Forebrain/Midbrain/Hindbrain - Narrow, Short, Truncated | | 1 | | | | | | | | | | 1 |
| Midbrain - Narrow | | | | | | | | | | | 1 | 1 |
| Midbrain - Not Evident | | | | | | | | | | 1 | | 1 |
| Midbrain - Short | | | | | | | | | | 1 | | 1 |
| Midbrain - Truncated | | | | | | | | | | 1 | 1 | 2 |
| **Pharyngeal Arches** | | | | | | | | | | | | |
| Not Remarkable | 15 | 26 | 9 | 12 | 10 | 8 | 12 | 9 | 15 | 22 | 15 | 153 |
| 1st Arch – Not Evident | | | | | | | | | | | 1 | 1 |
| 2nd Arch – Not Evident | | | | | | | | | | | 1 | 1 |
| **Rotation** | | | | | | | | | | | | |
| Not Remarkable | 11 | 21 | 8 | 9 | 8 | 3 | 10 | 9 | 13 | 15 | 10 | 117 |
| Bend | 3 | 4 | 1 | 2 | 2 | 4 | 1 | | 1 | 6 | 5 | 29 |
| Squirrel | | | | 1 | | | | | | | | 1 |
| S-Shaped/Turning | 1 | 1 | | | | 1 | 1 | | 1 | 1 | 1 | 7 |
| **Somites** | | | | | | | | | | | | |
| Not Remarkable | 14 | 26 | 9 | 12 | 10 | 8 | 11 | 9 | 15 | 22 | 16 | 152 |
| Fused | | | | | | | 1 | | | | | 1 |

| | | | | | | | | | | | | |
|---|---|---|---|---|---|---|---|---|---|---|---|---|
| Not Able to Visualize | 1 | | | | | | | | | | | 1 |
| Not well Defined | 1 | | | | | | | | | | | 1 |
| **Yolk Sac** | | | | | | | | | | | | |
| Not Remarkable | 11 | 24 | 9 | 11 | 9 | 7 | 12 | 8 | 14 | 20 | 16 | 141 |
| Abnormal Vasculature | 1 | | | 1 | 1 | | | | | | | 3 |
| Collapsed | 2 | 1 | | | | | | 1 | | 1 | | 5 |
| Embryo Outside Yolk Sac | | 3 | | | | 1 | 2 | 1 | 1 | 2 | | 10 |
| No Primary vessels | 1 | | | | 1 | | | | | | | 2 |
| Pale | 1 | | | | | | | | 1 | | | 2 |
| Visceral Yolk Sac Not Closed | | 1 | | | | 1 | | | 1 | 2 | | 5 |

*Data includes studies from the past 4 years

### Supplemental Table 4.

| Rabbit WEC Morphology Detailed Observations | | | | | |
|---|---|---|---|---|---|
| **Dose Level** | **0 µM** | **1 µM** | **10 µM** | **50 µM** | **100 µM** |
| **Able to Examine** | | | | | |
| Yes | 25 | 12 | 15 | 13 | 15 |
| No | 5 | 4 | 1 | 3 | 1 |
| Dead | 0 | 1 | 0 | 0 | 0 |
| Grossly Malformed | 0 | 0 | 0 | 1 | 1 |
| Not in bottle | 0 | 0 | 0 | 0 | 0 |
| Embryo Outside of Yolk Sac | 5 | 3 | 1 | 2 | 0 |
| **Caudal** | | | | | |
| Not remarkable | 24 | 10 | 13 | 12 | 13 |
| Caudal Dysgenesis | 0 | 0 | 0 | 0 | 0 |
| Other | 0 | 0 | 0 | 0 | 1 |
| Tail - Narrow | 0 | 0 | 0 | 0 | 0 |
| Tail – Remnant | 0 | 0 | 0 | 0 | 0 |
| Tail - Short; mild | 0 | 1 | 1 | 0 | 1 |
| Tail - Short; moderate | 1 | 1 | 1 | 1 | 0 |
| Total Embryos with Caudal Abnormalities | 1 | 2 | 2 | 1 | 2 |
| **Craniofacial** | | | | | |
| Not remarkable | 22 | 10 | 13 | 12 | 10 |
| Blister | 0 | 0 | 0 | 0 | 0 |
| Cleft Between Eye and First Arch | 0 | 0 | 2 | 0 | 0 |
| Lens - Not Evident | 0 | 0 | 0 | 0 | 0 |
| Loss Cells Below Eye or Between Eye and Arch | 0 | 0 | 1 | 0 | 0 |
| Mesencephalic Flexure - Misshapen/Malformed | 0 | 0 | 0 | 0 | 0 |
| Mesencephalic Flexure - Not Evident | 1 | 0 | 0 | 0 | 0 |
| Mesencephalic Flexure - Small/Narrow | 0 | 0 | 0 | 0 | 0 |
| Nasal Prominence - Not Evident | 0 | 0 | 0 | 0 | 0 |
| Nasal Prominence - Short; Moderate | 2 | 1 | 0 | 1 | 1 |
| Nasal Prominence - Short; Severe | 1 | 0 | 1 | 0 | 0 |
| Optic Vesicle - Loss of Definition | 0 | 0 | 0 | 0 | 3 |
| Optic Vesicle - Misshapen | 0 | 0 | 0 | 0 | 0 |
| Optic Vesicle - Not Evident | 0 | 0 | 0 | 0 | 0 |
| Optic Vesicle - Remnant | 1 | 0 | 1 | 0 | 0 |
| Optic Vesicle - Small | 0 | 0 | 0 | 0 | 0 |
| Other | 0 | 1 | 0 | 0 | 0 |
| Otic Placode - Displaced | 0 | 0 | 0 | 0 | 0 |
| Otic Placode - Loss of Definition | 0 | 1 | 0 | 0 | 0 |
| Otic Placode - Misshapen | 0 | 0 | 0 | 1 | 0 |

| | | | | | |
|---|---|---|---|---|---|
| Otic Placode - Not Evident | 0 | 0 | 0 | 0 | 0 |
| Otic Placode - Small | 0 | 0 | 1 | 0 | 0 |
| Maxillary process extends to 1st arch | 0 | 0 | 1 | 0 | 0 |
| Optic Vesicle - Round without lens | 0 | 0 | 0 | 0 | 1 |
| Maxillary process shorter than 1st arch | 1 | 0 | 0 | 0 | 0 |
| Olfactory - small indentations | 1 | 0 | 0 | 0 | 0 |
| Otic Placode - Round | 0 | 0 | 0 | 0 | 0 |
| Total Embryos with Craniofacial Abnormalities | 3 | 2 | 2 | 1 | 5 |
| **Forelimb Bud** | | | | | |
| Not remarkable | 17 | 8 | 11 | 7 | 10 |
| Not Evident | 1 | 1 | 0 | 3 | 0 |
| Other | 0 | 0 | 0 | 0 | 0 |
| Remnant | 0 | 1 | 2 | 0 | 3 |
| Small | 8 | 2 | 2 | 4 | 2 |
| Apical ectodermal ridge - Not evident | 0 | 0 | 0 | 0 | 0 |
| Total Embryos with Forelimb Bud Abnormalities | 8 | 4 | 4 | 6 | 5 |
| **Hindlimb Bud** | | | | | |
| Not remarkable | 22 | 10 | 13 | 11 | 9 |
| Not Evident | 0 | 2 | 2 | 1 | 0 |
| Other | 0 | 0 | 0 | 0 | 0 |
| Remnant | 2 | 0 | 0 | 0 | 4 |
| Small | 1 | 0 | 0 | 1 | 2 |
| Apical ectodermal ridge - Not evident | 0 | 0 | 0 | 0 | 0 |
| Total Embryos with Hindlimb Bud Abnormalities | 3 | 2 | 2 | 2 | 6 |
| **Heart** | | | | | |
| Not remarkable | 25 | 12 | 13 | 13 | 12 |
| Atrium - Small/Compressed | 0 | 0 | 0 | 0 | 0 |
| Atrium - Swollen | 0 | 0 | 0 | 0 | 0 |
| Cardiac Chambers - Cell Death | 0 | 0 | 0 | 0 | 0 |
| Chambers not well Defined | 0 | 0 | 0 | 0 | 0 |
| Damaged | 0 | 0 | 0 | 0 | 0 |
| Dextrocardia | 0 | 0 | 0 | 0 | 1 |
| No Heartbeat | 0 | 0 | 0 | 0 | 0 |
| Outflow tract - Abnormal Looping | 0 | 0 | 1 | 0 | 2 |
| Outflow tract - Kinked | 0 | 0 | 0 | 0 | 0 |
| Outflow tract - Narrow | 0 | 0 | 0 | 0 | 0 |
| Outflow tract - Not Evident | 0 | 0 | 0 | 0 | 0 |
| Outflow tract - Not Well Defined | 0 | 0 | 0 | 0 | 0 |
| Outflow tract - Swollen | 0 | 0 | 0 | 0 | 0 |
| Other | 0 | 0 | 0 | 0 | 0 |
| Pericardial Sac - Blebbing | 0 | 0 | 0 | 0 | 0 |

| | | | | | |
|---|---|---|---|---|---|
| Pericardial Sac - Cell Death | 0 | 0 | 0 | 0 | 0 |
| Pericardial Sac - Filled with Blood | 0 | 0 | 0 | 0 | 0 |
| Pericardial Sac – Not Evident | 0 | 0 | 0 | 0 | 0 |
| Pericardial Sac - Swollen | 0 | 0 | 1 | 0 | 1 |
| Tube-shaped | 0 | 0 | 0 | 0 | 0 |
| Ventricle - Small/Compressed | 0 | 0 | 0 | 0 | 0 |
| Ventricle - Swollen | 0 | 0 | 0 | 0 | 0 |
| 2 chambers | 0 | 0 | 0 | 0 | 0 |
| Total Embryos with Heart Abnormalities | 0 | 0 | 2 | 0 | 3 |
| **Neural Tube** | | | | | |
| Not remarkable | 22 | 11 | 13 | 11 | 11 |
| 4th Ventricle - Collapsed | 0 | 0 | 0 | 0 | 0 |
| 4th Ventricle - Swollen | 2 | 0 | 0 | 0 | 0 |
| Blister | 1 | 0 | 0 | 2 | 1 |
| Brain - Hemorrhage | 0 | 0 | 0 | 0 | 0 |
| Exencephaly | 0 | 0 | 0 | 0 | 0 |
| Forebrain - Cleft | 0 | 0 | 0 | 0 | 0 |
| Forebrain - Kinked | 0 | 0 | 0 | 0 | 0 |
| Forebrain - Misshaped | 0 | 0 | 0 | 0 | 0 |
| Forebrain - Narrow | 0 | 0 | 0 | 0 | 0 |
| Forebrain - Not Evident | 0 | 0 | 0 | 0 | 0 |
| Forebrain - Short | 1 | 0 | 1 | 0 | 0 |
| Forebrain - Truncated | 1 | 0 | 1 | 0 | 0 |
| Forebrain, Midbrain - Loss of Definition | 0 | 0 | 0 | 0 | 0 |
| Forebrain, Midbrain, Hindbrain - Loss of Definition | 1 | 1 | 0 | 0 | 0 |
| Forebrain, Midbrain, Hindbrain - Narrow | 0 | 0 | 0 | 0 | 0 |
| Forebrain, Midbrain, Hindbrain - Narrow, Short, Truncated | 1 | 0 | 1 | 1 | 0 |
| Forebrain, Midbrain, Hindbrain - Short | 0 | 0 | 0 | 0 | 0 |
| Forebrain, Midbrain, Hindbrain - Truncated | 0 | 0 | 0 | 0 | 0 |
| Hindbrain - Kinked | 0 | 0 | 0 | 0 | 0 |
| Hindbrain - Narrow | 0 | 0 | 0 | 0 | 0 |
| Hindbrain - Short | 0 | 0 | 0 | 0 | 0 |
| Hindbrain - Truncated | 1 | 0 | 0 | 0 | 0 |
| Midbrain - Indentation | 0 | 0 | 0 | 0 | 0 |
| Midbrain - Kinked | 0 | 0 | 0 | 0 | 0 |
| Midbrain - Narrow | 0 | 0 | 0 | 0 | 0 |
| Midbrain - Not Evident | 0 | 0 | 0 | 0 | 0 |
| Midbrain - Short | 1 | 0 | 0 | 0 | 1 |
| Midbrain - Truncated | 0 | 0 | 0 | 0 | 2 |
| Midbrain/Hindbrain - Loss of Definition | 0 | 0 | 0 | 0 | 0 |
| Open Neuropore - Caudal | 0 | 0 | 0 | 0 | 0 |

| | | | | | |
|---|---|---|---|---|---|
| Open Neuropore - Forebrain | 0 | 0 | 0 | 0 | 0 |
| Open Neuropore - Hindbrain | 0 | 0 | 0 | 0 | 0 |
| Other | 0 | 0 | 0 | 0 | 0 |
| Spinal Cord - Kinking | 0 | 0 | 0 | 0 | 0 |
| Spinal Cord - Lack of Neural Fold Fusion | 0 | 0 | 0 | 0 | 0 |
| Spinal Cord - Narrow | 0 | 0 | 0 | 0 | 0 |
| Spinal Cord - Swelling | 0 | 0 | 0 | 0 | 1 |
| Hindbrain - Pontine Flexure Dorsal aspect concave/sharp apex | 0 | 0 | 0 | 0 | 0 |
| Cell death | 1 | 1 | 1 | 1 | 1 |
| Midbrain - Suture Line Not Evident | 0 | 0 | 0 | 0 | 0 |
| Total Embryos with Neural Tube Abnormalities | 3 | 1 | 2 | 2 | 4 |
| **Pharyngeal Arches** | | | | | |
| Not remarkable | 25 | 11 | 14 | 12 | 15 |
| 1st Arch - Cell Death | 0 | 1 | 0 | 0 | 0 |
| 1st Arch - Hemorrhage | 0 | 0 | 0 | 0 | 0 |
| 1st Arch - Narrow/Short | 0 | 0 | 0 | 0 | 0 |
| 1st Arch – Not evident | 0 | 0 | 0 | 0 | 0 |
| 1st Arch - Remnant | 0 | 0 | 0 | 0 | 0 |
| 1st Arch - Swollen | 0 | 0 | 0 | 0 | 0 |
| 2nd Arch - Cell Death | 0 | 1 | 0 | 0 | 0 |
| 2nd Arch - Hemorrhage | 0 | 0 | 0 | 0 | 0 |
| 2nd Arch - Narrow/Short | 0 | 0 | 1 | 0 | 0 |
| 2nd Arch – Not evident | 0 | 0 | 0 | 0 | 0 |
| 2nd Arch - Remnant | 0 | 0 | 0 | 0 | 0 |
| Fused - 1st and 2nd | 0 | 0 | 0 | 0 | 0 |
| Fused - 1st at midline | 0 | 0 | 0 | 0 | 0 |
| Other | 0 | 0 | 0 | 0 | 0 |
| Ramus - Blister | 0 | 0 | 0 | 0 | 0 |
| Ramus - Cell Death | 0 | 0 | 0 | 1 | 0 |
| 3 Arches | 0 | 0 | 0 | 0 | 0 |
| 4 Arches | 0 | 0 | 0 | 0 | 0 |
| 3rd & 4th Arch – Not evident | 0 | 1 | 1 | 0 | 0 |
| Mandibular Process Not Fused | 0 | 0 | 0 | 0 | 0 |
| Total Embryos with Pharyngeal Arch Abnormalities | 0 | 1 | 1 | 1 | 0 |
| **Rotation** | | | | | |
| Not remarkable | 20 | 9 | 12 | 10 | 8 |
| Bend | 2 | 2 | 1 | 2 | 6 |
| Other | 0 | 0 | 0 | 0 | 0 |
| Squirrel | 2 | 1 | 2 | 1 | 0 |
| Squirrel - Tail Fused to Body | 0 | 0 | 0 | 0 | 0 |
| S-Shaped | 0 | 0 | 0 | 0 | 0 |

| | | | | | |
|---|---|---|---|---|---|
| S-Shaped/Turning | 1 | 0 | 0 | 0 | 1 |
| Total Embryos with Rotation Abnormalities | 5 | 3 | 3 | 3 | 7 |
| **Somite Morphology** | | | | | |
| Not remarkable | 25 | 11 | 14 | 13 | 12 |
| Cobblestone | 0 | 0 | 0 | 0 | 0 |
| Compressed | 0 | 0 | 0 | 0 | 0 |
| Fused | 0 | 0 | 0 | 0 | 0 |
| Missing | 0 | 0 | 0 | 0 | 0 |
| Narrow | 0 | 0 | 0 | 0 | 0 |
| Not Able to Visualize | 0 | 1 | 1 | 0 | 2 |
| Not well Defined | 0 | 0 | 0 | 0 | 1 |
| Other | 0 | 0 | 0 | 0 | 1 |
| Rounded | 0 | 0 | 0 | 0 | 0 |
| Serrated | 0 | 0 | 0 | 0 | 0 |
| Small | 0 | 0 | 0 | 0 | 0 |
| Split | 0 | 0 | 0 | 0 | 0 |
| Wedge Shaped | 0 | 0 | 0 | 0 | 0 |
| Total Embryos with Somite Abnormalities | 0 | 1 | 1 | 0 | 3 |
| **Yolk Sac Morphology** | | | | | |
| Not remarkable | 21 | 11 | 14 | 13 | 10 |
| Abnormal Vasculature | 2 | 1 | 1 | 0 | 1 |
| Blebbing/Blister | 0 | 0 | 0 | 0 | 2 |
| Blood Cells - Not Evident | 0 | 0 | 0 | 0 | 0 |
| Blood Cells - Sparse | 0 | 0 | 0 | 0 | 0 |
| Blood Island(s) | 1 | 0 | 0 | 0 | 1 |
| Collapsed | 2 | 0 | 0 | 0 | 1 |
| Discolored | 0 | 0 | 0 | 0 | 0 |
| Ectoplacental cone - Detached | 0 | 0 | 0 | 0 | 0 |
| Ectoplacental cone - Small | 0 | 0 | 0 | 0 | 0 |
| Invaginated | 0 | 0 | 0 | 0 | 0 |
| Necrotic | 0 | 0 | 0 | 0 | 0 |
| Not Examined - Hole in Visceral yolk sac | 0 | 0 | 0 | 0 | 0 |
| Other | 0 | 0 | 0 | 0 | 0 |
| Pale | 0 | 0 | 1 | 0 | 2 |
| Rough Surface | 0 | 0 | 0 | 0 | 0 |
| Visceral yolk sac Not Closed | 0 | 0 | 0 | 0 | 0 |
| No Primary vessels | 0 | 0 | 0 | 0 | 0 |
| Total Yolk Sacs with any abnormalities | 4 | 1 | 1 | 0 | 5 |

*Dextrocardia observations were recorded in the raw data, but not summarized, because dextrocardia is considered a culture-related artifact and not a test item effect.
